# Sensory adaptation and pupil-linked arousal support flexible evidence accumulation during perceptual decision making

**DOI:** 10.64898/2026.02.03.703553

**Authors:** Kara D. McGaughey, Joshua I. Gold

**Affiliations:** Department of Neuroscience, University of Pennsylvania, Philadelphia, PA 19104; Computational Neuroscience Initiative, University of Pennsylvania, Philadelphia, PA 19104; Neuroscience Graduate Group, University of Pennsylvania, Philadelphia, PA 19104

## Abstract

Effective decision making in dynamic environments requires flexible evidence accumulation. Although models often express this flexibility as a property of the accumulator, its implementation in the brain may involve adaptive mechanisms operating at other stages of the decision process. We examined two such mechanisms: 1) stimulus-specific sensory adaptation at the level of evidence encoding; and 2) arousal-related neuromodulation, which could, in principle, affect both evidence encoding and accumulation. We measured single-unit activity in the middle temporal area (MT) and pupil-linked arousal while monkeys performed a modified random-dot motion direction-discrimination task in which an adapting stimulus with varied temporal stability preceded a behaviorally relevant test stimulus. The monkeys’ decisions reflected adaptive evidence accumulation that depended on temporal-context stability and corresponded to context-dependent changes in both stimulus-specific sensory adaptation in MT and task-evoked pupil responses. However, adaptation and pupil adjustments were not related to each other. Together, these findings suggest that multiple mechanisms contribute to flexible, context-dependent evidence accumulation, including changes in sensory adaptation that shape evidence encoding and changes in arousal that may shape the accumulation process itself.

## Introduction

Perceptual decisions often require the accumulation of uncertain sensory evidence over time. This accumulation process is complicated by the fact that the nature and source of incoming evidence can change even while accumulation is underway. Under these changing conditions, effective decision makers should adjust how sensory information is accumulated so that their perceptual judgments reflect current conditions rather than outdated evidence gathered before a change occurred (Gold & Stocker, 2017). This flexibility can be achieved by adjusting the evidence-accumulation process in several ways, including adopting a leak and/or saturating non-linearity that can be adjusted according to environmental conditions to balance sensitivity to stability versus change (Glaze et al., 2015; Usher & McClelland, 2001). These adaptive forms of evidence accumulation are key features of decision-making behavior in dynamic environments (Glaze et al., 2015; Murphy et al., 2021; Piet et al., 2018) and are often modeled using one or more parameters that control the temporal dynamics of an accumulator (Glaze et al., 2015; Usher & McClelland, 2001; Veliz-Cuba et al., 2016, Fig. 1).

**Fig. 1:**
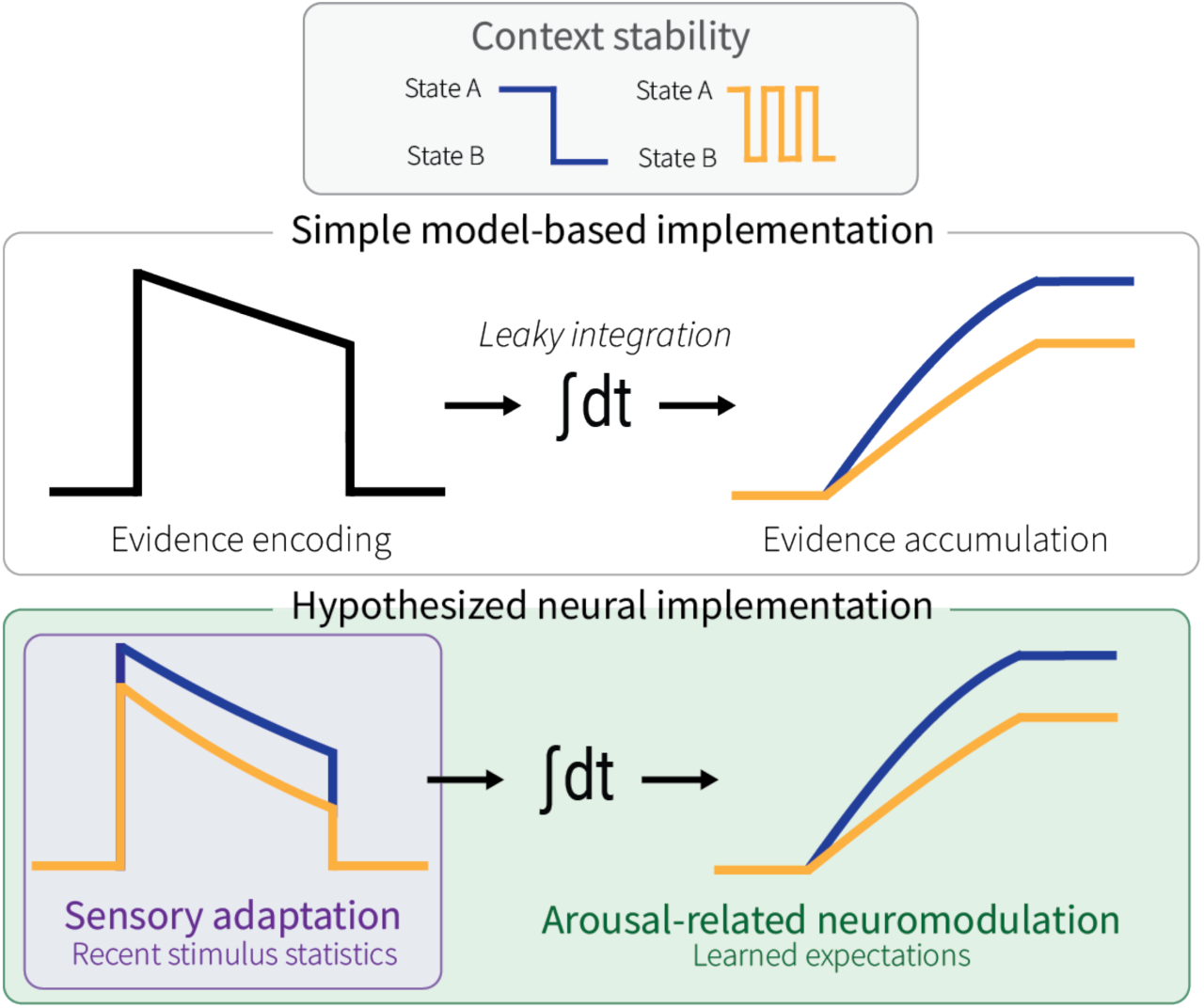
Hypothesized neural implementation of context-dependent changes in evidence accumulation. For perceptual decisions about motion direction, momentary evidence is encoded in the middle temporal area (MT) and then accumulated by downstream circuits to form a decision variable that guides behavior. Models of decision-making often implement flexible, context-dependent evidence accumulation via a single parameter that controls the temporal dynamics (e.g., leakiness) of an accumulator (*top*). In principle, adjustments to the accumulation process could involve changes in evidence encoding, accumulation, or both (*bottom*). Here we used manipulations of context stability (low-versus high-frequency direction reversals of an adapting motion stimulus) to test for a role of evidence encoding. Because global signals related to arousal can influence decision flexibility and cortical information processing at multiple levels, we further considered potential interactions between adaptation- and arousal-related effects on flexible evidence accumulation.

However, unlike these simple models, the brain contains many mechanisms that could, in principle, shape the temporal dynamics of evidence accumulation. Here, we considered a potential role for stimulus-specific sensory adaptation of individual neurons encoding the evidence that is accumulated downstream to form the decision (Fig. 1). Adaptation is a hallmark of sensory systems, occurring across modalities (Dalton, 2000; Lampl & Katz, 2017; Laughlin, 1989; Ulanovsky et al., 2004) and species (Brenner et al., 2000; Clarke et al., 2015; Lesica et al., 2007; Nagel & Doupe, 2006; Tolias et al., 2001). Recent theoretical work has framed adaptation as a form of active inference, whereby sensory systems not only modulate their responses to match the statistical structure of incoming signals, but also infer when those statistics have changed and how rapidly they are expected to vary (DeWeese & Zador, 1998; Młynarski & Hermundstad, 2021; Solomon & Kohn, 2014; Wark et al., 2007; Weber et al., 2019). Consistent with this idea, recordings from isolated retinal ganglion cells have shown that the temporal dynamics of recent stimulus history can modulate adaptation (Wark et al., 2009). Yet it remains unknown whether individual neurons in primate sensory cortex undergo similar temporal context-dependent adaptation, and whether such adaptation contributes to flexible evidence accumulation for perceptual decision making.

To test these ideas, we leveraged decisions about visual motion. These decisions depend on neurons in the middle temporal area (MT) of extrastriate visual cortex (K. Britten et al., 1992; K. H. Britten et al., 1996; Hanks et al., 2006; Newsome & Pare, 1988; Salzman et al., 1990), which exhibit robust stimulus-specific sensory adaptation (Kohn & Movshon, 2003; Van Wezel & Britten, 2002). We manipulated the stability of recent sensory experience by modifying the classic random-dot motion direction-discrimination task to include extended exposure to an adapting stimulus that reversed direction at either a low or high frequency, creating two context-stability conditions (Fig. 2A). Immediately following this adapting stimulus, monkeys viewed and reported the direction of a test stimulus, which was identical across conditions, allowing us to isolate the influence of recent stimulus statistics on both behavior and neural activity. We found that monkeys adjusted their decision-making behavior in a manner that was consistent with context-dependent changes in the temporal dynamics of evidence accumulation. These adjustments were based, in part, on differences in how the evidence being accumulated was encoded by individual MT neurons.

**Fig. 2:**
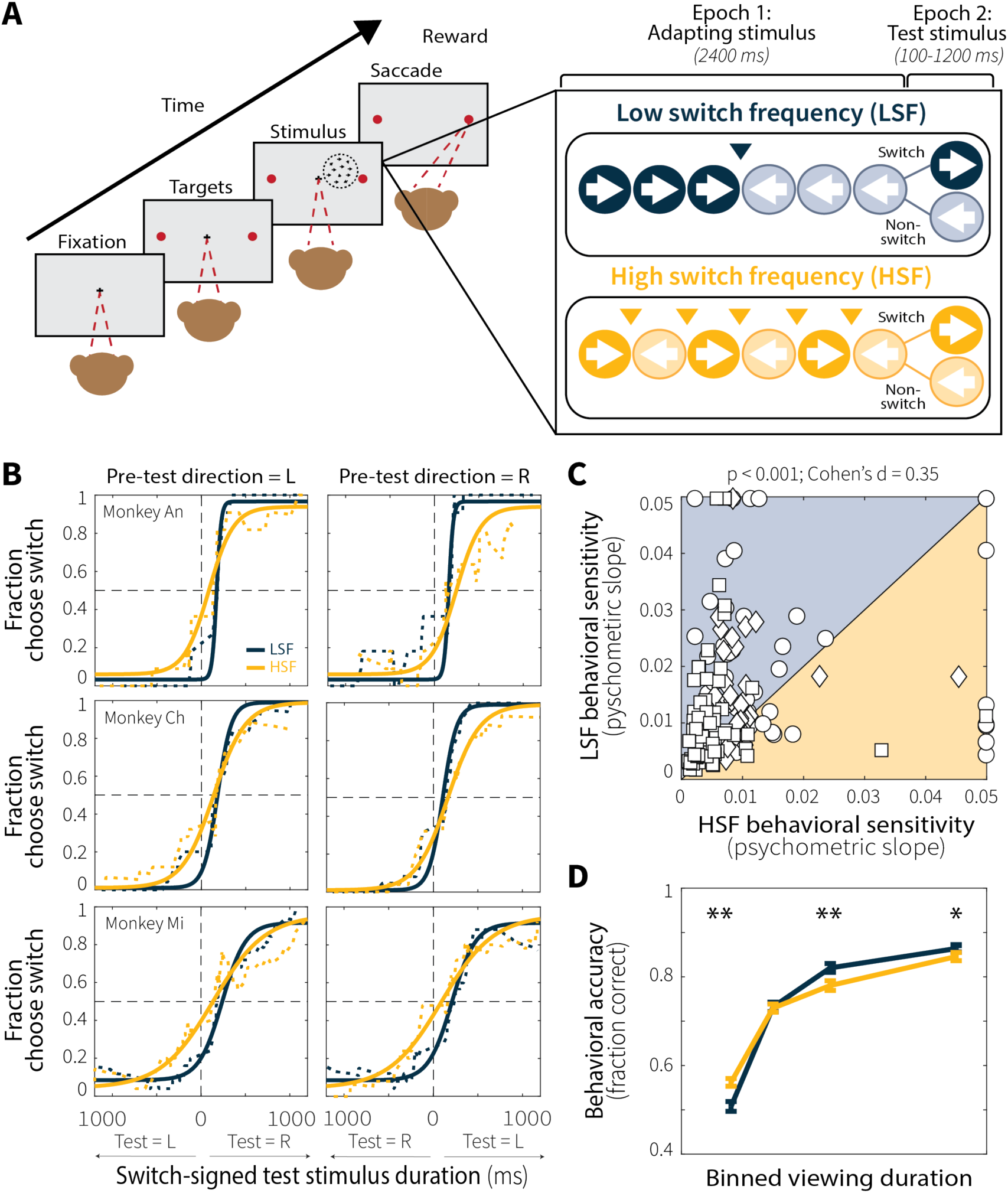
Manipulating context stability in a random-dot motion task affects evidence-accumulation behavior. **(A)** Each trial consisted of an adapting stimulus (2400 ms) followed immediately by a test stimulus (100–1200 ms, drawn from an exponential distribution). During the adapting stimulus, random-dot motion switched directions at either a low (LSF; 1 switch) or high (HSF; 5 switches) frequency, creating two context-stability conditions. Between epochs, there was a 50% probability of an additional change in motion direction, producing switch and non-switch trials. After presentation of the test stimulus, the fixation point was extinguished and the monkey reported the final direction of motion with a saccade to the corresponding choice target. **(B)** Running average (square window size=5 trials, sorted by test-stimulus duration) of behavioral choice data (dotted line) and psychometric fits (solid lines) for representative sessions from each monkey. LSF (blue) and HSF (orange) conditions were fit separately. Shallower slopes indicate decreased perceptual sensitivity as a function of viewing time. **(C)** Across-session comparison of fitted slopes for LSF and HSF conditions (Monkey An = 51 sessions, circles; Ch = 34, diamonds; Mi = 76, squares). *p-*value is from a Wilcoxon signed-rank test for equal medians. (**D**) Behavioral performance on switch trials as a function of test-stimulus duration, which was binned (100–225 ms, 225–375 ms, 375–600 ms, and 600–1200 ms) to include roughly equal numbers of trials (mean ± SEM across sessions). Asterisks denote bins for which LSF and HSF performance differed significantly (Wilcoxon signed-rank test for equal medians; * *p* < 0.05; ** *p* < 0.001).

We also examined whether and how these context-dependent behavioral and neural adjustments related to simultaneous measures of pupil diameter. Non-luminance-mediated fluctuations in pupil diameter, an index of arousal, track quantities relevant to adaptive evidence accumulation, including environmental volatility and temporal expectations (Akdoğan et al., 2016; Nassar et al., 2012; Schwiedrzik & Sudmann, 2020). Pupil diameter also relates to perceptual sensitivity, bias, and variability in evidence accumulation (de Gee et al., 2014, 2017; Keung et al., 2019; Krishnamurthy et al., 2017; Murphy et al., 2014). These findings imply a role for the arousal system in optimizing perceptual decision making, likely via neuromodulatory effects on neural information-processing dynamics (Aston-Jones & Cohen, 2005; Devilbiss et al., 2006; Joshi & Gold, 2020; Navarra et al., 2017; Waterhouse et al., 1998; Wyrick & Mazzucato, 2021). Arousal systems are therefore well positioned to, in principle, influence how sensory information is encoded, accumulated, or both (Fig. 1). Our results, which show a dissociation between sensory adaptation-related and arousal-related effects on evidence accumulation, provide new insights into the complex, distributed mechanisms that contribute to flexible decision making.

## Results

We trained three rhesus macaques (An, Ch, and Mi) to perform a version of a random-dot motion direction-discrimination task that combined reversals in dot-motion direction (Glaze et al., 2015; Fig. 2A) with elements of an adaptation-test paradigm (Van Wezel & Britten, 2002). Each trial consisted of two sequential epochs: an adapting epoch (Epoch 1) and a test epoch (Epoch 2). During the adapting epoch (2400 ms), the random-dot motion stimulus abruptly switched direction at either a low (LSF; 1 switch, occurring at 1200 ms) or high (HSF; 5 switches, occurring every 400 ms) frequency. The two adapting (“context-stability”) conditions were designed such that the total exposure time to each adapting motion direction was matched for low and high switch frequencies (1200 ms per direction). Immediately following the adapting epoch, the test stimulus was shown for a duration that was sampled independently on each trial from a truncated exponential distribution (100–1200 ms), requiring the monkeys to accumulate evidence over varied and unpredictable timescales. At the end of this duration, the test stimulus and fixation point were extinguished simultaneously, cueing the monkeys to report the final motion direction by making a saccade to the corresponding choice target.

Critically, for both low and high switch-frequency conditions there was a 50% probability that the motion stimulus changed direction between the adapting and test epochs. This design ensured that switch and non-switch trials were equally likely, and thus that the adapting stimulus provided no information about the correct test-stimulus direction. Moreover, the transition between adapting and test epochs was marked by predictable timing and a reduction in motion coherence (to ensure that the task was difficult enough to require sustained evidence accumulation), implying that the monkeys could, in principle, learn to ignore the adapting stimulus altogether. Instead, as detailed below, systematic effects of context stability on behavior, neural activity, and pupil modulations point to inherent ways in which the brain uses recent stimulus history to shape perception.

### Context stability modulates evidence-accumulation behavior

To quantify if and how the monkeys’ decisions depended on recent temporal context, we fit their choice behavior using a time-dependent logistic function (Equation 1) that predicted the probability of reporting a switch in motion direction as a function of test-stimulus viewing time and trial type (switch or non-switch; Fig. 2B). This model captured behavior well, with Tjur’s pseudo-R² values that exceeded those computed by shuffling the association between test-stimulus durations and switch/non-switch trial types (Extended Data Fig. 1).

The adapting stimulus affected the monkeys’ decisions in several ways. On the shortest-duration switch trials, monkeys tended to make choices congruent with the final direction of the adapting stimulus, reflected in lower accuracy and a rightward shift of the psychometric functions (examples shown in Fig. 2B; mean ± SEM shift across sessions = 168.25 ± 8.38 ms for Monkey An, 110.79 ± 9.76 ms for Monkey Ch, 138.46 ± 10.35 ms for Monkey Mi). It was not until after ∼140 ms that their choices began to more strongly reflect the direction of the test stimulus. Together, these effects indicate that the monkeys were not ignoring the adapting stimulus but instead were using at least some of its directional information to inform their decisions, particularly when given only brief exposure to the test stimulus. As the duration of the test stimulus increased, performance accuracy also tended to increase, including more “switch” choices on switch trials, suggesting that the monkeys’ decisions became more strongly dependent on the test as opposed to the adapting stimulus.

Critically, despite identical test-stimulus properties, durations, and equal probabilities of a behaviorally relevant switch, the rate of increase in accuracy as a function of viewing time differed between context-stability conditions. Psychometric slopes were shallower for the high relative to low switch-frequency condition (Fig. 2B–C; individual animals in Extended Data Fig. 2), reflecting diminished sensitivity to incoming evidence as a function of viewing time. As a consequence, performance on switch trials showed a “crossover” dynamic, such that accuracy was higher in the high relative to low switch-frequency condition at short durations but higher in the low relative to high switch-frequency condition at long durations (Fig. 2D; pairwise comparison in Extended Data Fig. 3A). This pattern of behavior is consistent with adaptive adjustments in evidence accumulation, in which a higher leak rate or lower saturating non-linearity facilitates performance on short switch trials by downweighting pre-change evidence but limits performance on longer-duration switch trials as more recent, relevant evidence is also discounted. These context-dependent temporal dynamics are inconsistent with a simple shift in the timing of the onset of evidence accumulation (either advanced or delayed relative to test-stimulus onset), which would shift the psychometric curves but not change their slopes. Instead, these slope changes show that recent temporal context can shape the temporal dynamics of decision formation.

### Context stability shapes sensory adaptation in MT

We recorded activity from 155 MT single units while the three monkeys performed the task (Monkey An = 55, Ch = 13, Mi = 87). As expected, given that stimulus properties were tailored to each neuron (see Methods), the activity of both individual units (Fig. 3A) and the recorded population (Fig. 3B) were robustly modulated by motion direction, showing strong responses to motion in each cell’s preferred direction and weak responses in the opposite, anti-preferred (“null”) direction that followed stimulus switching dynamics. Given the lack of responsiveness to null motion and the unbalanced exposure to the final motion direction on non-switch trials (i.e., an additional 800 ms at low relative to high switch frequency), unless otherwise noted the analyses of MT neural activity were restricted to preferred-motion switch trials, which were most informative for characterizing context-dependent evidence encoding.

**Fig. 3:**
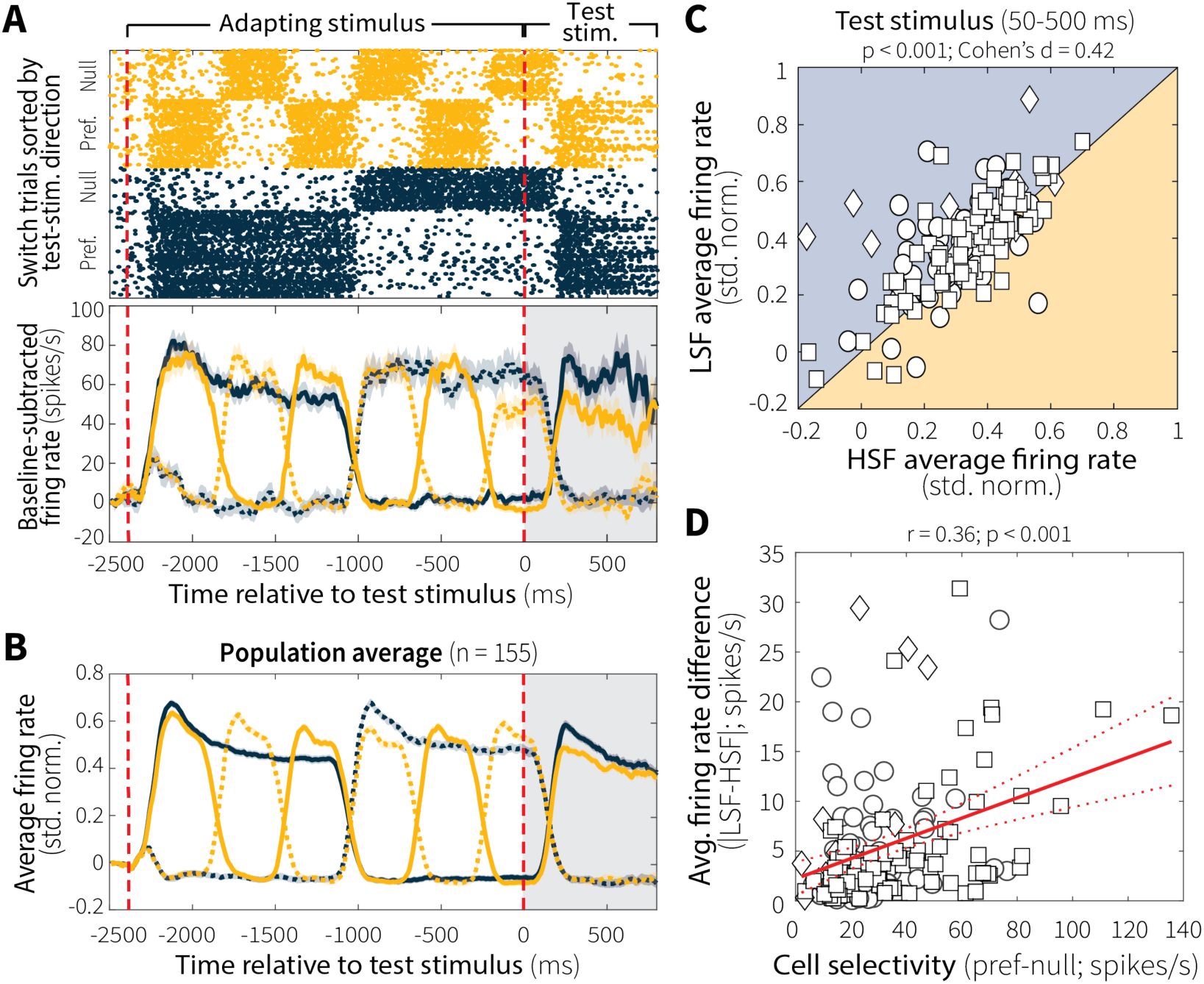
Context stability modulates motion-evidence encoding in MT. (**A**) Raster plot (top) and baseline-subtracted average firing rate (bottom; mean ± SEM) for switch trials from a representative MT single unit on low (LSF; blue) and high (HSF; orange) switch-frequency trials. Solid and dashed lines indicate responses to motion in the unit’s preferred and null directions, respectively, defined relative to motion direction during the test stimulus. (**B**) Baseline-subtracted, normalized firing rate averaged across all recorded MT single units (*n* = 155; mean ± SEM). (**C**) Pairwise comparison of average single-unit activity during the test stimulus (50–500 ms) for LSF and HSF preferred-motion switch trials (Monkey An = 55 single units, circles; Ch = 13, diamonds; Mi = 87, squares). *p-*value is from a Wilcoxon signed-rank test for equal medians. (**D**) Pearson’s correlation between MT direction selectivity and the magnitude of LSF–HSF response differences during the test stimulus (50–500 ms). Red solid and dashed lines indicate a linear fit ± 95% CI.

Temporal-context stability systematically affected responses to preferred motion during the test epoch, with reduced activity following an adaptor that switched at a high versus low frequency (Fig. 3C; individual animals in Extended Data Fig. 4A). Differences in neural activity between the two context-stability conditions were best characterized by a change in overall response magnitude rather than a change in temporal dynamics (Extended Data Fig. 4B). Moreover, these differences varied systematically with direction selectivity. MT single units with stronger direction selectivity, defined as larger differences between responses to preferred and null motion, showed greater differences in activity between switch-frequency conditions (Fig. 3D). In other words, neurons that encoded motion evidence more strongly and more selectively were also more sensitive to the temporal statistics and stability of that evidence, exhibiting larger context stability-dependent adjustments.

These context-dependent differences in MT neural activity did not reflect baseline differences in responsiveness that persisted throughout the low and high switch-frequency blocks. Rather, they emerged over the course of each trial, consistent with stimulus-specific sensory adaptation. Specifically, the average population responses to the first presentation of preferred motion during the adapting stimulus were initially matched between conditions (Fig. 4A; although one animal showed an early difference, possibly reflecting learned expectation about switching dynamics within blocks, Extended Data Fig. 5). Context-dependent differences developed with repeated stimulus presentation within a trial, culminating in robust differences in neural responses to the test stimulus between low and high switch-frequency conditions (Fig. 4A,B)

**Fig. 4:**
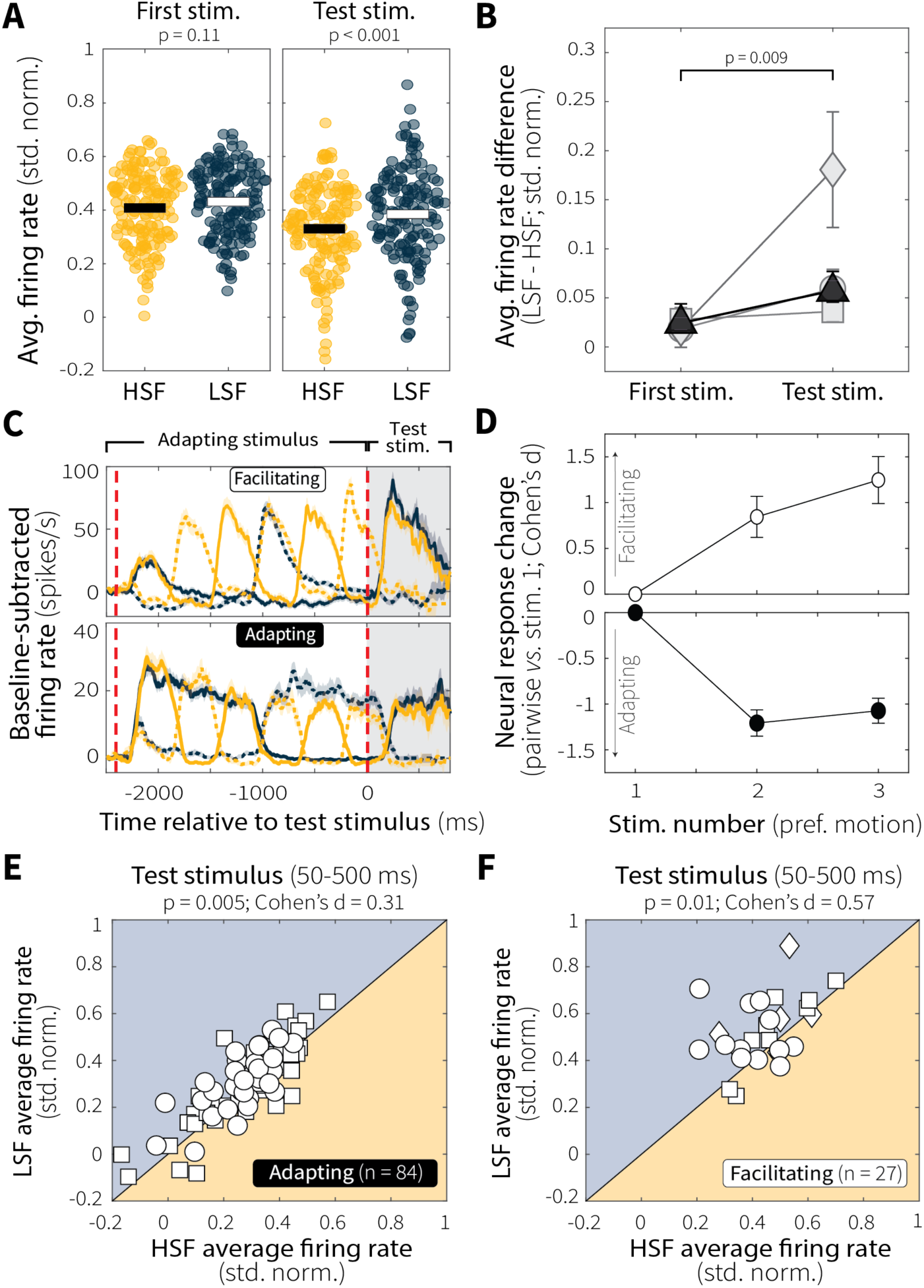
Context-dependent differences in MT evidence encoding emerge following repeated preferred-motion stimulus presentation. (**A**) Responses of MT single units (points) to the first (adapting-stimulus onset) and final (test) presentations of preferred-motion stimuli on switch trials. Horizontal bars represent means. *p-*values are for two-sample *t*-tests for equal means of the two distributions. (**B**) Change in response differences (LSF–HSF) from the first (adapting-stimulus onset) and final (test) presentations of preferred-motion stimuli on switch trials. Individual animals are in gray; group average is in black; all points are mean ± SEM across units. *p-*value is from a two-sample *t*-test comparing the full distribution of response differences (LSF–HSF) from the two time intervals for all monkeys. (**C**) Example switch-trials responses from representative facilitating (top) and adapting (bottom) single units. Colors and line styles as in Fig. 2A. (**D**) Comparison of average baseline-subtracted and normalized responses across successive presentations of preferred motion within the adapting epoch for HSF switch trials (panel C, orange solid). For adapting (black, n = 87) and facilitating (white, n = 27) groups, responses to each preferred-motion presentation were compared to the first (adapting-stimulus onset) preferred-motion response using Cohen’s d to quantify changes in response magnitude as a function of stimulus number. Points denote mean ± SEM across units. (**E**, **F**) Pairwise comparison of average single-unit responses to the test stimulus (50–500 ms) on switch trials during LSF versus HSF conditions for adapting (**E**) and facilitating (**F**) units. Different symbols represent data from different monkeys, as in Fig. 2C. *p-*values are from a Wilcoxon signed-rank test for equal medians.

The effect of context stability on evidence encoding was consistent across distinct patterns of adaptation. Whereas most MT single units showed persistent response decrements with repeated preferred-motion presentations (Fig. 4C, *bottom*), some exhibited response increments (Fig. 4C, *top*). We classified these two groups of neurons as “adapting” and “facilitating,” respectively, based on changes in baseline-subtracted, normalized responses between the first presentation of preferred motion and preferred motion during the test stimulus (see Methods). Regardless of whether neurons were adapting or facilitating, both groups showed comparable magnitudes of response change as a function of preferred-motion stimulus presentation (Fig. 4D). Notably, the majority of this response change, particularly for adapting neurons, occurred between the first and second presentations of preferred motion, with relatively little additional change across subsequent presentations. This pattern suggests that adaptation dynamics are shaped by temporal features of the adapting stimulus, such as the frequency of direction switches, as opposed to the number of preferred-motion stimulus presentations. Moreover, regardless of whether MT neurons were adapting or facilitating, both groups exhibited robust context-stability effects with diminished activity at high versus low switch frequency during the test stimulus (Fig. 4E,F). Thus, context-specific differences in evidence encoding arose from stimulus-specific sensory adaptation that unfolded over repeated preferred-motion exposure and was expressed across MT neurons with different response profiles.

### Context-dependent evidence encoding in MT relates to evidence-accumulation behavior

To assess the behavioral relevance of context-dependent differences in MT neural responses, we first quantified how well individual neurons discriminated between preferred and null motion at low versus high switch frequency using ROC area (Fig. 5A). Across the population, ROC area tended to be lower for high versus low switch frequency (Fig. 5B; individual animals in Extended Data Fig. 6A), indicating decreased evidence discriminability in MT when the test stimulus was preceded by a more unstable adapting context. We then tested whether, across sessions, these differences in ROC area predicted context-dependent behavioral differences. Because ROC analyses were restricted to switch trials (which, unlike non-switch trials, have balanced exposure to the final adapting motion direction), we also restricted the behavioral analyses to switch trials, using percent correct instead of psychometric slope (which was based on all trials). We focused on trials 375–600 ms in duration, because that time bin exhibited robust context-dependent differences in accuracy that remained reliably above chance, but below ceiling, for both conditions (Fig. 2D and Extended Data Fig. 3B). We found that the differences in MT evidence discriminability (LSF minus HSF ROC area, in the epoch 200–400 ms following test-stimulus onset that showed the largest context-dependent differences in neural activity) correlated with context-stability differences in behavioral performance (LSF minus HSF percent correct for trials ending 375–600 ms following test-stimulus onset, Fig. 5C), such that greater differences in discriminability between low and high switch-frequency conditions predicted larger differences in performance. This effect was significant across animals, but driven primarily by Monkey An (Extended Data Fig. 6B).

**Fig 5:**
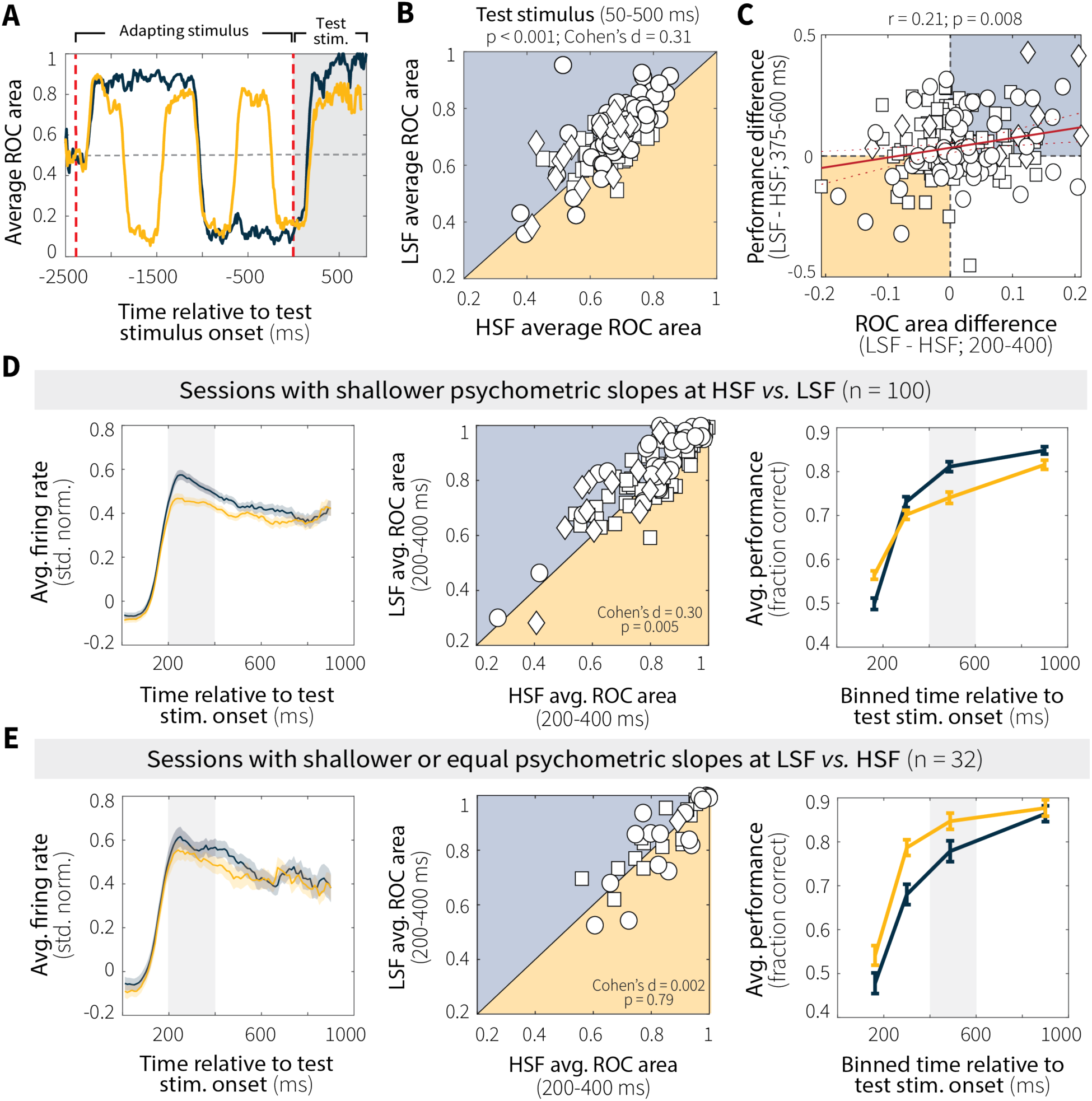
MT neural activity relates to evidence-accumulation behavior. (**A**) Average ROC area over time for a representative MT single unit showing discriminability between preferred and null motion during low (LSF, blue) and high (HSF, orange) switch-frequency switch trials. (**B**) Pairwise comparison of average ROC area during the test stimulus (50–500 ms) for LSF and HSF switch trials. *p-*value is from a Wilcoxon signed-rank test for equal medians. (**C**) Context stability-dependent differences in behavioral performance differences (LSF–HSF, measured as the difference in accuracy for trials that ended 375–600 ms after test-stimulus onset; gray band in D and E, left panels) plotted as a function of MT single-unit discriminability (LSF–HSF ROC area, 200–400 ms after test-stimulus onset; gray band in D and E, right panels). Red solid and dashed lines indicate a linear fit ± 95% CI. *p*-value is for the Pearson’s correlation coefficient (*H_0_*: *r* = 0). (**D**) Sessions with shallower psychometric slopes at HSF relative to LSF. Columns show (left) MT evidence encoding (mean ± SEM baseline-subtracted, normalized firing rates averaged across units), (middle) MT evidence discriminability (average ROC area, 200–400 ms after test-stimulus onset; *p*-values from a Wilcoxon signed-rank test for equal medians), and (right) behavioral performance in stimulus-duration bins with roughly equal numbers of switch trials (mean ± SEM across sessions). (**E**) Same as D, but for sessions with shallower or equal slopes at LSF relative to HSF. Different symbols in the four scatterplots represent data from different monkeys, as in Fig. 2C.

To further explore this relationship, we divided sessions based on evidence-accumulation behavior. For sessions in which the monkeys had shallower psychometric slopes at high relative to low switch frequency (Fig. 5D), MT neurons exhibited a corresponding decrease in firing rate and discriminability on switch trials (pairwise comparisons for firing rates in Extended Data Fig. 7). Conversely, for all other sessions (Fig. 5E), we did not identify any reliable context-stability differences in MT evidence encoding or discriminability (pairwise comparisons for firing rates in Extended Data Fig. 7). This relationship was specific to correct trials and absent for incorrect trials, which showed no modulation based on evidence-accumulation behavior (Extended Data Fig. 8). A similar, behavior-dependent pattern in discriminability was also evident on non-switch trials (Extended Data Fig. 9). However, these context-dependent differences were restricted to a smaller and earlier time window (50–200 ms), likely reflecting activity driven primarily by the adapting rather than the test stimulus. Together, these results suggest that adaptation-driven differences in MT evidence encoding contribute, at least in part, to context-dependent evidence-accumulation behavior.

### Context-dependent evoked pupil modulations relate to evidence-accumulation behavior

To probe whether processes beyond local sensory adaptation contribute to adaptive evidence accumulation, we examined both baseline and evoked pupil diameter as proxies for arousal-related modulations that have been shown to encode key features of adaptive perceptual decision making (Cheadle et al., 2014; Krishnamurthy et al., 2017; Murphy et al., 2021). Trial-wise baseline pupil diameter was correlated with the time required for monkeys to obtain fixation (Extended Data Fig. 10A), a relationship that was robust across sessions and animals (Extended Data Fig. 10B). However, baseline pupil diameter did not differ systematically between low and high switch-frequency conditions (Extended Data Fig. 10C), suggesting that it captured global as opposed to context-dependent fluctuations in arousal or task engagement. We therefore focused on task-evoked, as opposed to baseline, pupil responses in subsequent analyses.

Evoked pupil responses were modulated by context stability. Following a brief, stereotyped constriction caused by the abrupt luminance increase at motion-stimulus onset, differences in pupil diameter emerged between low and high switch-frequency conditions (Fig. 6A). To quantify these context-dependent effects, we used sliding-window linear regressions (Equation 2) to estimate differences in evoked pupil diameter (LSF–HSF), while controlling for baseline pupil diameter (because evoked and baseline diameter tend to be inversely related; Mathôt et al., 2018). For both monkeys (Monkey Ch was excluded from pupil-related analyses; see Methods), the effect of context-stability on evoked pupil diameter increased over the course of the adapting stimulus (Fig. 6B; individual animals in Extended Data Fig. 11A). This result is consistent with the idea that evoked pupil responses track context-dependent expectations or anticipatory processes associated with behaviorally relevant stimuli.

**Fig. 6:**
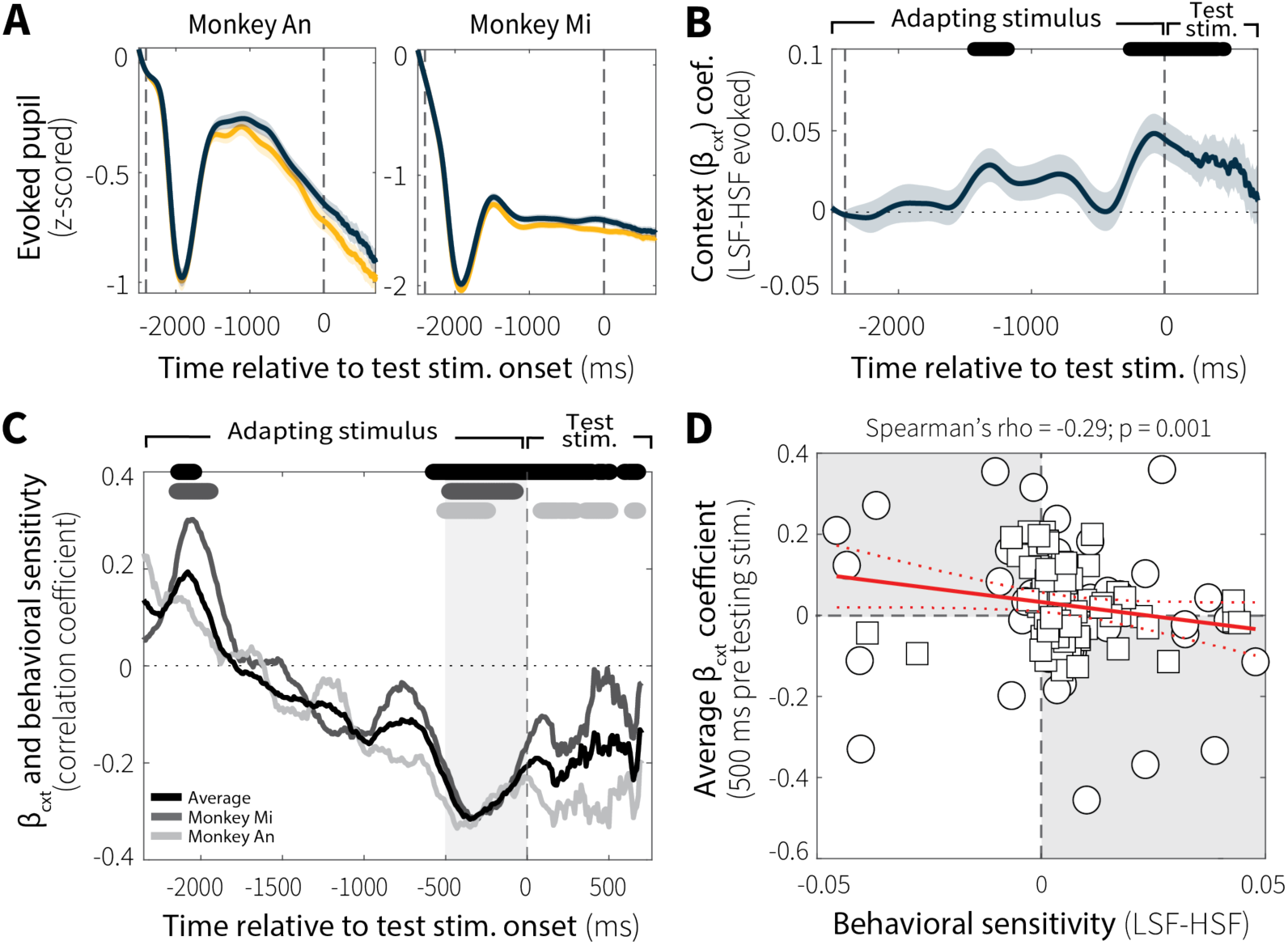
Evoked pupil responses depend on context stability and relate to evidence-accumulation behavior. (**A**) Average evoked pupil responses over time (z-scored per session) for low (LSF; blue) and high (HSF; orange) switch-frequency conditions (mean ± SEM across all sessions), shown separately for the two monkeys. (**B**) Sliding-window regression coefficients (β_cxt_; mean ± SEM across switch and non-switch trials for all sessions from both monkeys) estimating the effect of context-stability condition (LSF–HSF) on pupil diameter during stimulus viewing while controlling for baseline pupil diameter. Black bar (top) indicates windows in which β_cxt_ differed significantly from zero (*p* < 0.05, uncorrected for multiple comparisons). (**C**) Spearman’s rank correlation coefficient between behavioral sensitivity (LSF–HSF psychometric slope) and β_cxt_ coefficients from individual sessions as a function of time relative to test-stimulus onset (computed in 100 ms bins with 10 ms steps). Data from individual animals are shown separately (Monkey An: light grey; Monkey Mi: dark grey) along with the across-animal average (black). Corresponding grayscale bars at the top indicate time points with significant correlations (*p* < 0.05, uncorrected for multiple comparisons). (**D**) Spearman’s rank correlation between average evoked pupil diameter differences (β_cxt_, −500–0 ms relative to test-stimulus onset; gray band in C) and behavioral sensitivity differences (LSF–HSF slope) across sessions. Red solid and dashed lines indicate a linear fit ± 95% CI. *p*-value is for the Spearman’s correlation coefficient (*H_0_*: *rho* = 0).

Context-dependent differences in evoked pupil diameter were relevant to behavior. Specifically, differences in evoked pupil diameter (LSF–HSF) showed correlations with behavioral sensitivity (LSF–HSF psychometric slope) that varied over the course of the trial and peaked just before the onset of the test stimulus (Fig. 6C). Around this peak (−500–0 ms before test-stimulus onset), average context-dependent differences in evoked pupil diameter reliably predicted subsequent differences in behavioral sensitivity across sessions (Fig. 6D; individual animals in Extended Data Fig. 11B). As such, sessions with larger differences in evoked pupil diameter also tended to show larger context-dependent differences in behavioral sensitivity.

### Adaptation and arousal-related mechanisms are jointly and differentially recruited across sessions

To directly compare the relative contributions of MT sensory adaptation and arousal-related mechanisms to adaptive evidence-accumulation behavior, we separately incorporated trial-wise measures of MT neural activity and evoked pupil diameter into the psychometric functions (Equation 3). This approach allowed us to quantify how strongly each signal modulated behavioral sensitivity as a function of context stability (LSF–HSF) on a session-by-session basis.

Both neural and pupil terms improved model fits, as evidenced by increased explanatory power relative to the behavior-only model in both animals (Fig. 7A). However, the relative improvement attributable to neural activity versus evoked pupil diameter differed between monkeys, with Monkey An showing a larger difference between neural versus pupil terms than Monkey Mi (Wilcoxon rank-sum test on neural–pupil differences in Tjur’s pseudo-R²; p < 0.001, Fig. 7A), implying joint, but flexible recruitment of adaptation and arousal-related mechanisms.

**Fig. 7:**
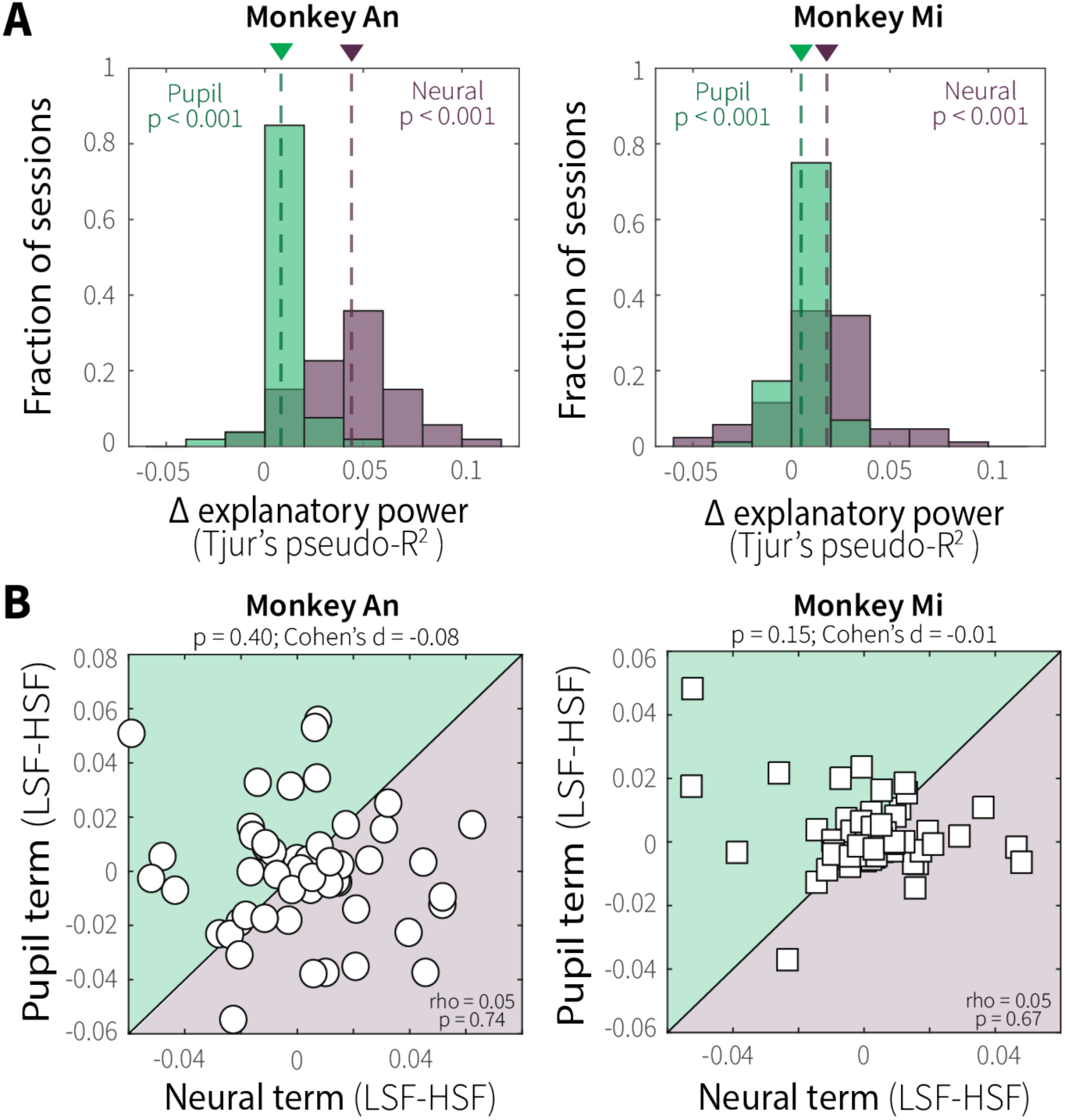
Context-stability jointly and differentially recruits adaptation and arousal-related mechanisms across sessions. (**A**) Distributions of differences in explanatory power (Tjur’s pseudo-R²) for models fit with versus without the neural (purple) or pupil (green) term across sessions for Monkey An (left) and Monkey Mi (right). Mean values are indicated by dashed lines and corresponding colored triangles. *p-*values are from a one-sample sign test for *H_0_*: fit parameters came from a distribution with a median of zero. (**B**) Context-stability differences (LSF–HSF) in neural (MT firing rate; abscissa) and pupil (evoked diameter; ordinate) model terms for Monkey An (left) and Monkey Mi (right). Points are fits to data from individual sessions/units from logistic models that quantified how much the slope of time-dependent psychometric functions covaried with the given neural or pupil term, computed separately for LSF and HSF trials. Shaded quadrants denote relative dominance of neural (purple) or pupil (green) effects. *p-*value*s* are from a Wilcoxon signed-rank test for equal medians (top) and Spearman’s correlation (bottom).

This flexible recruitment was evident across sessions, where the influence of context-dependent MT neural activity and evoked pupil diameter on behavioral sensitivity varied considerably, with neither term reliably predominating over the other (Fig. 7B). Session-wise, context-dependent contributions from neural and pupil terms were uncorrelated with each other, consistent with the two signals contributing separately rather than redundantly or in opposition (Fig. 7B). In further support of this idea, context-dependent modulations of pupil size were not related systematically to context-dependent modulations of MT neural activity or discriminability (Extended Data Fig. 11C,D). Together, these findings suggest that sensory adaptation and pupil-linked arousal both contribute to adaptive evidence accumulation, likely operating at different processing stages, with their relative weighting varying substantially across sessions and animals.

## Discussion

Flexible evidence accumulation is critical for effective decision making in dynamic environments. Many algorithmic accounts propose that this flexibility arises from context-dependent adjustments to the process of evidence accumulation itself. However, the brain contains many adaptive mechanisms at different stages of the decision process that could contribute to this behavioral flexibility. Motivated by this idea, we manipulated the temporal stability of recent sensory experience and simultaneously measured both neural activity in the middle temporal area (MT) and pupil-linked arousal. We identified two complementary mechanisms that could support flexible evidence accumulation, one involving sensory adaptation and the other involving arousal-related neuromodulation (Fig. 1). Our results suggest that these mechanisms operate separately, likely at different processing stages, providing a new view of the distributed neural substrates for flexible evidence accumulation in dynamic environments.

Our MT findings indicate that context-dependent changes in how choice accuracy evolves as a function of viewing time, which is a behavioral signature of flexible evidence accumulation, can arise, at least in part, from context-dependent differences in sensory encoding. Both motion-evidence encoding and discriminability of MT responses during the test-stimulus tended to be reduced following exposure to more temporally unstable adapting stimuli (Fig. 3A-C). These reductions were not sustained differences throughout the two context-stability conditions but instead emerged on each trial through repeated presentations of preferred motion at different switch frequencies, consistent with stimulus-specific sensory adaptation (Fig. 4A-B). This stimulus-specific adaptation varied systematically across individual units, such that more direction-selective cells were more affected by differences in stimulus temporal statistics, which is consistent with previous findings that more active MT neurons adapt more strongly (Fig. 3D; Van Wezel & Britten, 2002).

It is well established that sensory adaptation can depend on stimulus statistics such as the mean, variance, and salience of sensory inputs (Dean et al., 2005; Fairhall et al., 2001) and even changes in those statistics (Kastner & Baccus, 2011; Tikidji-Hamburyan et al., 2015). Here, we extend these findings by demonstrating that adaptation can also depend on the rate of change of the stimulus statistics. This kind of temporal dynamic-dependent adaptation has been observed in retinal ganglion cells (Wark et al., 2009). Our work extends those findings to non-human primate cortical neurons with causal links to behavior (Hanks et al., 2006; Newsome & Pare, 1988), in particular showing that context stability-dependent adaptation in MT shapes the sensory evidence available for time-evolving perceptual decisions.

We also observed heterogeneity in how MT neural activity changed over the course of the adapting stimulus, with some cells exhibiting progressive response attenuation and others facilitation (Fig. 4C-D). A plausible mechanism for this facilitation is that adaptation alters the local excitation-inhibition balance (Higley & Contreras, 2007; Solomon & Kohn, 2014). In particular, repeated presentation of null motion may weaken opponent input, producing disinhibition of preferred-motion responses (Kohn & Movshon, 2003). Consistent with this account, facilitating neurons showed increased responses to preferred motion following null-motion adaptation (Extended Data Fig. 12). This diversity of adaptation profiles mirrors observations in retinal ganglion cells (Kastner & Baccus, 2011, 2013), primary auditory cortex (Seay et al., 2020), and somatosensory cortex (Adibi et al., 2013; Cohen-Kashi Malina et al., 2013; Derdikman et al., 2006; Kheradpezhouh et al., 2017), and recent work suggests that whether a neuron adapts or facilitates may itself be a flexible property that depends on recent sensory experience (Dobler et al., 2024). Nevertheless, regardless of the adaptation profile, context stability shaped MT activity and discriminability in ways that corresponded to context-dependent differences in evidence-accumulation behavior across sessions.

These adaptation-driven changes in MT evidence encoding were relevant to behavior but not fully prescriptive of context-dependent adjustments in evidence accumulation. For instance, for sessions in which the monkeys showed behavioral evidence of context-dependent evidence-accumulation (i.e., reduced sensitivity as a function of viewing time at high relative to low switch frequency; Fig. 5D), there was a corresponding change in MT evidence encoding (e.g., reduced neural activity at high relative to low switch frequency; Fig. 5D). However, sessions showing the opposite pattern of behavior did not show opposite neural effects (Fig. 5E). Thus, while sensory adaptation supports flexible evidence accumulation by shaping the fidelity of sensory input, additional mechanisms likely contribute to how that information is dynamically weighed and accumulated over time.

Consistent with this idea, fluctuations in task-evoked pupil size, an index of arousal-related neuromodulation, were also related systematically to temporal-context stability. Differences in evoked pupil diameter emerged between low and high switch-frequency conditions over the course of the adapting epoch, peaking just before test-stimulus onset (Fig. 6B). This time course suggests that evoked pupil dynamics may be tracking expectations about environmental statistics to prepare downstream decision circuits for the behaviorally relevant stimulus, consistent with previous work (Filipowicz et al., 2020). Across sessions, the magnitude of context-dependent differences in evoked pupil diameter predicted corresponding differences in behavioral sensitivity (Fig. 6C-D). Specifically, larger pupil diameter corresponded to less-efficient evidence accumulation, a result consistent with previous work relating larger pupil diameter to increased variability (Murphy et al., 2014) and individual differences in the temporal dynamics of evidence accumulation (Keung et al., 2019). These findings join a growing body of work showing that pupil-linked arousal contributes to adaptive mechanisms that shape behavior to match current environmental demands.

Our findings further suggest that these pupil-linked arousal mechanisms operate separately from sensory adaptation to support flexible evidence accumulation. Adding trial-wise MT neural activity or evoked pupil diameter each increased explanatory power relative to behavior-only models, demonstrating that both signals provided behaviorally relevant information (Fig. 7). However, the fact that their relative contributions were uncorrelated across sessions and that evoked pupil responses were unrelated to MT activity suggests that sensory adaptation and pupil-linked arousal may shape behavior through distinct mechanisms at different processing stages. That said, given the general unreliability of trial-wise data, particularly pupil responses, we cannot rule out the possibility that the lack of correlation between adaptation and arousal-related contributions results, in part, from this inherent variability as opposed to true independence of the underlying mechanisms.

One possible interpretation of these arousal-related signals is that they are related to neuromodulatory mechanisms that control the efficiency or leakiness of downstream accumulator circuits, such as those in the lateral intraparietal area (LIP), frontal eye field (FEF), or superior colliculus (SC), that read out MT activity. Alternatively, or in addition, both MT adaptation and pupil-linked arousal could be independently regulated by prefrontal regions, such as the dorsolateral prefrontal cortex (dlPFC), anterior cingulate cortex (ACC), or orbitofrontal cortex (OFC), that monitor task performance and environmental state (Behrens et al., 2007; McGuire et al., 2014; O’Reilly et al., 2013) and project both to brainstem arousal systems (Breton-Provencher & Sur, 2019; Schwarz & Luo, 2015) and to sensory cortex (Banerjee et al., 2020; Gilbert & Li, 2013; Murphy et al., 2021). Either possibility supports a framework in which sensory adaptation provides a “bottom-up” mechanism that modulates evidence encoding based on recent stimulus statistics, whereas pupil-linked arousal provides a “top-down” signal that adjusts how that encoded evidence is accumulated over time according to expectations about environmental stability. Together, these complementary processes may jointly implement the context-dependent nonlinearities described by normative models of evidence accumulation in dynamic environments (Glaze et al., 2015; Murphy et al., 2021; Fig. 1), with variability in their relative contributions across sessions and animals suggesting flexible recruitment that may depend on factors such as training, experience, or strategy.

Our conclusions are tempered by several limitations. First, we do not know the exact algorithmic nature of the context-dependent effects we measured on evidence accumulation behavior (i.e., differences in how accuracy evolves as a function of viewing time), which could involve a leak, saturating non-linearity, and/or other form of adjustment. Disentangling these alternatives necessitates additional behavioral testing and analyses (e.g., reverse-correlation analyses, which would require interspersing lower-coherence test stimuli than were used in this study) under a broader range of conditions. Second, we do not know where the context-dependent adaptation measured in MT originates. In principle, these effects could, at least in part, be inherited from upstream visual areas, including areas V1 (Born & Bradley, 2005; Churchland et al., 2005; Movshon & Newsome, 1996; Rodman et al., 1989) and V2 (DeYoe & Van Essen, 1985), where adaptation to motion is also known to occur. Third, our recordings sampled single units across sessions and thus did not capture population-level changes in MT activity. Adaptation can influence activity beyond single neurons, including changes in correlated variability within neural populations that can affect decision making (Solomon & Kohn, 2014; Whitmire & Stanley, 2016). Moreover, correlated variability in cortex can covary with pupil size and LC-NE activity (Joshi & Gold, 2022; Vinck et al., 2015), providing another potential link between arousal and sensory encoding that merits further study. Fourth, downstream cortical (Ding & Gold, 2012; Kim & Shadlen, 1999; Shadlen & Newsome, 2001) and subcortical (Ding & Gold, 2013; Horwitz & Newsome, 1999; Lovejoy & Krauzlis, 2010) areas that read out MT activity to form the decision may also contribute to flexible adjustments to the process of evidence accumulation. Future work combining further behavioral testing and simultaneous recordings and/or causal manipulations across sensory, decision, and arousal-related regions will be essential to disentangle how multiple distributed mechanisms implement adaptive evidence accumulation to support flexible decision making.

In summary, this work contributes to a broader reconceptualization of classic, hierarchical views of sensorimotor pathways that cast primary sensory cortex as a static encoder supplying input to flexible downstream decision processes that guide behavior. Instead, a growing body of evidence suggests that these early sensory areas can provide adaptive, context-dependent signals shaped by recent experience and environmental structure that can impact behavior (Waiblinger et al., 2022, 2025). Our demonstration of separable, behaviorally relevant contributions from sensory adaptation in MT and pupil-linked arousal supports this view, indicating that flexible decision making in dynamic environments emerges from distributed mechanisms that include intrinsically adaptive signals in early sensory cortex that complement adjustments in downstream processes.

## Materials and Methods

Three adult male rhesus monkeys (*Macca mulatta*) participated in this study. Two of the monkeys (An and Ch) had extensive prior training on a standard (i.e., non-switching) version of the random-dot motion direction-discrimination task. All training, surgery, and experimental procedures conformed to the National Institutes of Health Guide for the Care and Use of Laboratory Animals and were approved by the University of Pennsylvania Institutional Animal Care and Use Committee (protocol #806027).

### Behavioral task

The behavioral task (Fig. 1A) combined a change-point variant of the random-dot motion task (Glaze et al., 2015) with adaptation-test elements from traditional MT sensory adaptation experiments (Van Wezel & Britten, 2002). Briefly, each trial consisted of two sequential epochs: an adapting-stimulus epoch (2400 ms) and a test-stimulus epoch (100–1200 ms, sampled independently from an exponential distribution with a mean of 500 ms). During the adapting epoch, a continuous random-dot motion stimulus was presented, within which motion direction switched at either a low (LSF; 1 switch) or high (HSF; 5 switches) frequency, creating two context-stability conditions. Change-point frequencies were designed so that total exposure time to each motion direction was matched across conditions (1200 ms per direction). Between epochs, there was a 50% probability of an additional change in motion direction, yielding an approximately equal number of switch and non-switch trials within each session. During the test epoch, the monkey maintained fixation and accumulated motion evidence over variable durations before reporting the final motion direction with a saccade to one of two choice targets. A juice reward was provided following correct choices.

For Monkey An (all 51 sessions), Mi (all 76 sessions), and Ch (11 out of 35 sessions), low (LSF) and high (HSF) switch-frequency conditions were run in blocks with counterbalanced order across sessions. The remaining sessions for Monkey Ch (n = 24) included interleaved LSF and HSF trials. Initial motion direction was counterbalanced across trials within each session. The adapting stimulus used high motion coherence (70%) to drive robust MT neural responses and adaptation, whereas the test stimulus used lower motion coherence (50–60%, adjusted across sessions to maintain overall task performance near ∼70–80% correct) to promote temporal integration.

Eye position was monitored with a video-based system (EyeLink, SR Research) sampled at 1000 Hz and used to enforce fixation during motion viewing, register the saccadic response, and provide reward/error feedback online based on comparisons between the saccade endpoint and target locations.

### Analysis of behavioral data

Behavioral choice data were fit with a logistic function that modeled the probability of reporting a switch in motion direction as a function of test-stimulus duration and trial type (switch versus non-switch):

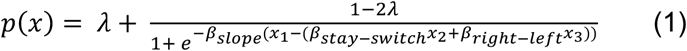

where *λ* is the error (“lapse”) rate independent of motion information (which here could arise for numerous reasons, including not just lapses of attention but also possibly inappropriate associations with motion directions observed during the adapting epoch); *β*_*slope*_ governs the steepness of the psychometric function as a function of test-stimulus viewing time (*x*_1_), signed by trial type (i.e., multiplied by 1 for switch trials and −1 for non-switch trials); *β*_*stay*−*s*w*itch*_ captures biases related to reporting switch versus non-switch (x_2_; switch = 1, stay = 0); and *β*_*ri*g*ht*−*left*_ captures biases related to right versus left (x_3_; right = 1, left = 0) choices. Here, “right” and “left” refer to all motion directions on the right or left side of vertical, respectively (e.g., motion directions between 270°–90° labeled “right,” and 90°–270° labeled “left”). Bias terms were encoded with respect to the x-intercept because the steep slopes can make the y-intercept difficult to estimate. We imposed an upper bound of 0.05 on *β*_*slope*_, which captured the majority of variability across sessions. This formulation allowed for the construction of a time-dependent psychometric function in which *β*_*slope*_ reflects evidence accumulation over time. Low and high switch-frequency conditions were fit separately to allow for comparison of fitted slope parameters.

### Electrophysiology

General surgical procedure and data acquisition methods have been described previous in detail (Law & Gold, 2008). Briefly, monkeys were prepared for experiments via surgical implantation of a head-holding device and recording cylinder. Area MT was targeted using stereotaxic coordinates, magnetic resonance imaging, and 3D reconstruction with BrainSight (Rogue Research, Inc). We searched for MT neurons with consistent spatial, direction, and speed tuning using a 99% coherence random-dot motion stimulus.

For each recording, multiple properties of the random-dot motion stimulus including aperture size and location, speed, and net motion direction were adjusted to match the tuning properties of the isolated single-unit and thus maximize stimulus-specific sensory adaptation. As such, exact stimulus parameters varied across sessions with stimulus sizes ranging from 4.6–15 degrees, locations within the contralateral visual hemifield ± 5 degrees from the point of fixation, speeds 2– 8 degrees/s, and motion-directions distributed roughly evenly within circular space. The adapting stimulus began with preferred or anti-preferred motion chosen at random with equal probability and counterbalanced across trials.

Neural activity was recorded using polyamide-coated tungsten electrodes (FHC, Inc) or glass-coated tungsten electrodes (Alpha-Omega). Spike waveforms were stored and sorted offline using software from Plexon, Inc. In some sessions, multiple MT single units were recorded simultaneously (n = 40 units in 19 sessions).

### Analysis of MT neural data

Firing rates were computed for each neuron and trial condition using a 100 ms sliding window advanced in 10 ms steps and aligned to the onset of the test stimulus. For each MT unit, firing rates were averaged across trial conditions, baseline subtracted on a trial-by-trial basis using responses from the 100 ms window immediately preceding adapting-stimulus onset as baseline, and normalized by the maximum firing rate observed during the 2400 ms adapting stimulus. For analyses requiring an average firing rate during the test stimulus, MT neural activity was averaged from a specified onset to either test-stimulus offset or the end of the specified time window, whichever came first. Only correct trials were included in analyses unless otherwise indicated.

Direction selectivity was quantified as the average difference between responses to preferred and anti-preferred (null) motion from 100 to 500 ms after adapting stimulus onset, corresponding to the overlapping stimulus period (400 ms) across switch-frequency conditions with a 100 ms delay.

To classify units as “adapting” versus “facilitating,” average baseline-subtracted and normalized neural activity was compared between the first presentation of preferred motion (200–400 ms following adapting stimulus onset) and preferred motion during the test stimulus (50–500 ms). Units were classified as adapting if responses decreased by >2.5% and as facilitating if responses increased by >2.5%, under both LSF and HSF conditions. We chose a liberal criterion for classification because we were interested in maintaining enough neurons in each group for statistical comparison of sensory adaptation as a function of preferred-motion stimulus presentation. Alternatively, using linear fits to peak preferred-motion responses and classifying neurons based on whether the fitted slope differed significantly from zero (*p* < 0.05), yielded 34 units (21.9%) with negative slopes classified as “adapting” and 4 units (0.3%) with positive slopes classified as “facilitating,” which are comparable to prior work in sensory cortical areas (Dobler et al., 2024; Kheradpezhouh et al., 2017; Seay et al., 2020).

### Analysis of pupil data

For each session, pupil data were aligned to the onset of the adapting stimulus and preprocessed by: (1) linear interpolation of missing values, (2) applying a first-order 5 Hz low-pass Butterworth filter, (3) removing and linearly interpolating samples exceeding ± 2 standard deviations from the mean, (4) subtracting the session-averaged response time course, and (5) z-scoring. For evoked-pupil analyses, only correct trials were included. For baseline pupil analyses, all trials were included so as not to disrupt the session’s temporal order.

Monkey Ch was excluded from pupil analysis because many sessions used interleaved (versus blocked) context-stability conditions, which removed the ability to establish trial-by-trial expectations of stability. Two sessions for Monkey An were also excluded because the pupil data were not properly saved.

To quantify context-stability effects on evoked-pupil dynamics, we applied sliding-window linear regressions (100 ms window, 10 ms slide; Equation 2) to estimate differences in evoked pupil response between LSF and HSF trials (*β*_2_) while controlling for baseline pupil (*β*_1_).

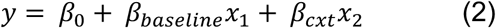

where, *β*_*baseline*_ estimates influence of baseline pupil diameter (*x*_1_: 100 ms window preceding motion stimulus onset) and *β*_*cxt*_ indexes context-stability condition (*x*_2_: 1 = LSF, 0 = HSF).

### Quantification of neural and pupil contributions to behavior

To quantify how neural and pupil measures contributed to behavioral sensitivity on a session-by-session basis, we added a term, *β_trial-wise_*, to our time-dependent logistic function (Equation 3).

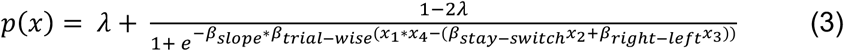

where, *β*_*trial*−w*ise*_ estimates the contribution of trial-wise MT activity for neural fits or trial-wise evoked pupil diameter for pupil fits (*x*_4_).

For these fits, MT activity was computed as the average firing rate during the test stimulus on each trial with a 50 ms delay. Because MT neurons exhibit minimal response to null motion, only trials ending in preferred motion were used for neural fits. Evoked pupil diameter was calculated as the average of the residuals (500 ms window preceding test-stimulus onset) from a linear fit with average pupil diameter and baseline pupil diameter. Otherwise, parameterization of the fits was as described in Equation 1. Given the lack of evidence for an interaction between pupil-linked arousal and sensory adaptation, neural and pupil terms were assessed as the sole contributor in separate fits. When assessing explanatory power, for each session, the reported pseudo-R^2^ value was the average across pseudo-R^2^ values from separate fits to low (LSF) and high (HSF) switch-frequency trials. Tjur’s pseudo-R² is a coefficient of discrimination. As such, its value increases only when additional model terms improve the separation of predicted probabilities across outcomes, providing a conservative basis for comparing models of differing complexity.

## Data and code availability

All data and code are available on Box at: https://upenn.box.com/s/7dxf9enb1e1nx19tszrqaur8wrtrflf4

## Acknowledgments

We thank Jean Zweigle for remarkable and diligent animal care as well as Jafar Bhatti, Long Ding, Catrina Hacker, Victoria Subritzky Katz, and Lowell Thompson for insightful comments on the manuscript. This work was supported by the NIH National Eye Institute (R01 EY015260; JIG), the NIH National Institute of Mental Health (R01 MH127566; JIG), and the National Science Foundation Graduate Research Fellowship Program (NSFGRFP DGE-1845298; KDM).

## Author contributions

Conceptualization, KDM and JIG; methodology, KDM and JIG; investigation, KDM; visualization, KDM and JIG; funding acquisition, KDM and JIG; project administration, KDM; supervision, JIG; writing – original draft, KDM; writing – review and editing, KDM and JIG.

---

**Extended Data Fig. 1:**
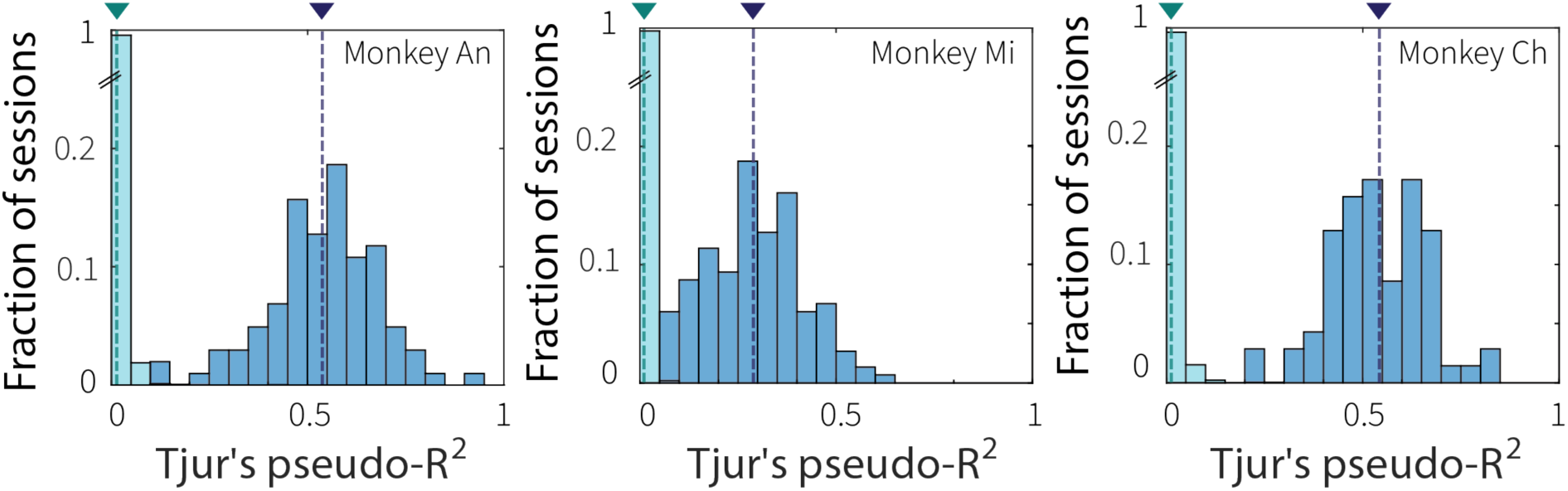
Time-dependent logistic model captures choice behavior. Distribution of Tjur’s pseudo-R^2^ values from empirical logistic fits (blue) compared with values from shuffled control (“null”) fits (cyan) for each monkey, as indicated. Null fits were obtained by shuffling the association between test-stimulus durations and switch/non-switch trial types across 100 iterations per session. For each session, the reported pseudo-R^2^ value was the average from separate fits to low and high switch-frequency trials. Dashed lines and triangles indicate mean values for each distribution. Enhanced explanatory power of the time-dependent logistic model versus the shuffled null was consistent across the three monkeys (Wilcoxon rank-sum test for equal medians: Monkey An: *p* < 0.001; Monkey Mi: *p* < 0.001; Monkey Ch: *p* < 0.001).

**Extended Data Fig. 2:**
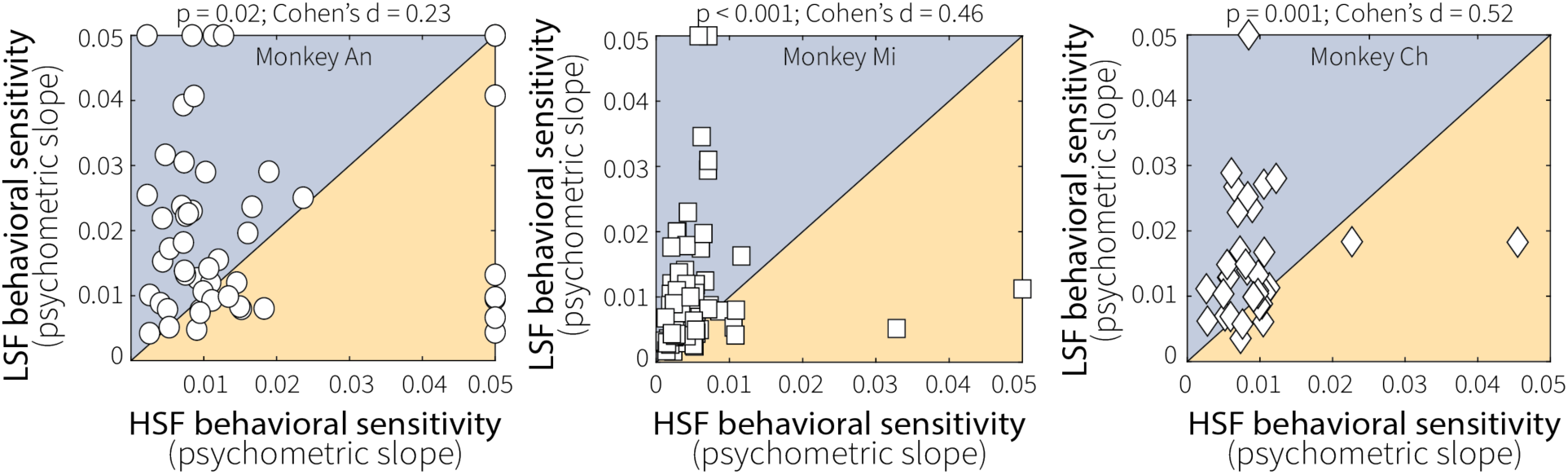
Context-dependent differences in evidence accumulation were consistent across animals. Pairwise comparisons of fitted psychometric slopes for low (LSF) and high (HSF) switch-frequency conditions across sessions for each monkey, as indicated. Slope values were bounded at 0.05, which is roughly the maximum resolvable steepness given our sampling of viewing times. All three monkeys tended to have shallower time-dependent psychometric slopes at HSF relative to LSF (*p*-values are from a Wilcoxon signed-rank test for equal medians). This result implies that the accuracy of the monkeys’ decisions improved more slowly as a function of viewing time at HSF relative to LSF.

**Extended Data Fig. 3:**
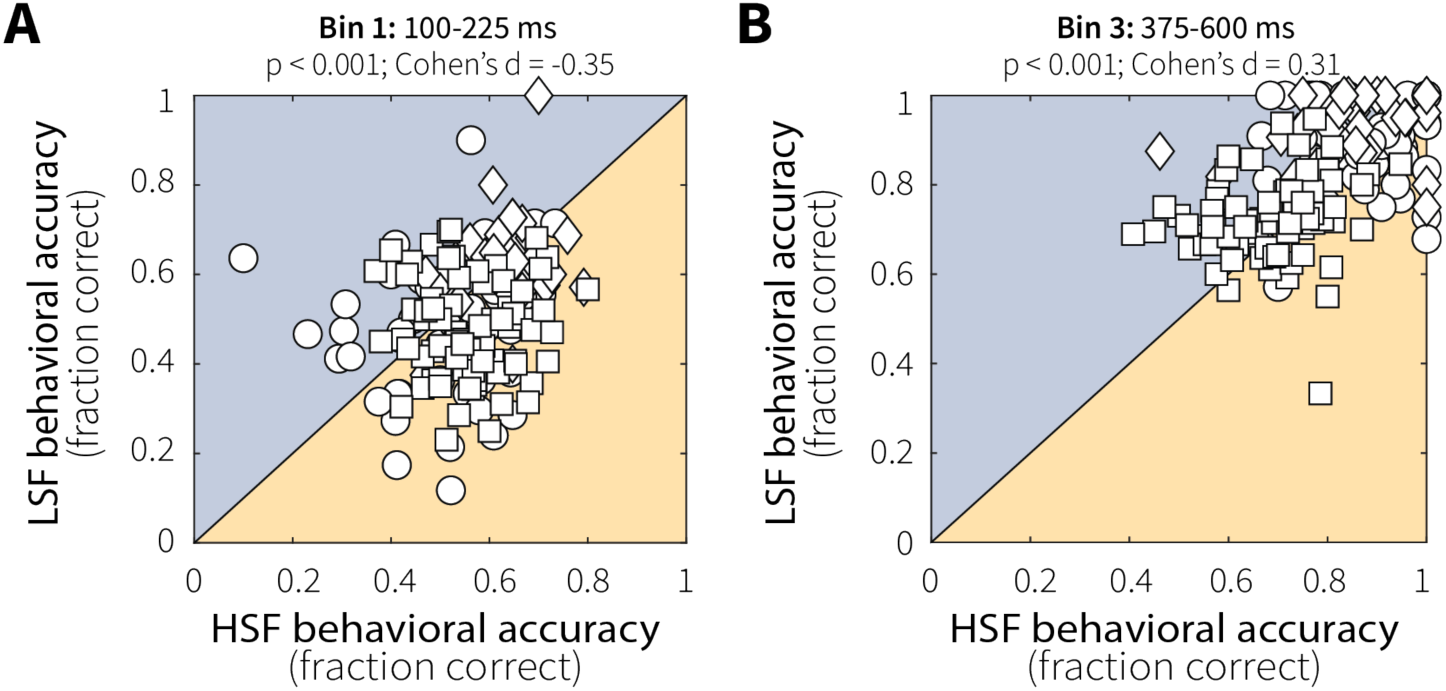
Differences in behavioral performance were consistent with context-dependent adjustments in evidence accumulation. Pairwise comparisons of behavioral accuracy (fraction correct) on switch trials for low (LSF) and high (HSF) switch-frequency conditions, shown separately for shorter-duration (A; Bin 1: 100– 225 ms) and longer-duration (B; Bin 3: 375–600 ms) test stimuli. On shorter-duration trials, accuracy was higher in the HSF relative to the LSF condition, whereas at longer viewing durations LSF accuracy surpassed HSF (*p*-values are from a Wilcoxon signed-rank test for equal medians). This crossover pattern is consistent with adaptive adjustments in evidence accumulation, in which a higher leak rate or lower saturating non-linearity facilitates performance on short switch trials by downweighting irrelevant pre-change evidence but limits performance on longer-duration switch trials as more recent, relevant evidence is also discounted (Glaze et al, 2015).

**Extended Data Fig. 4:**
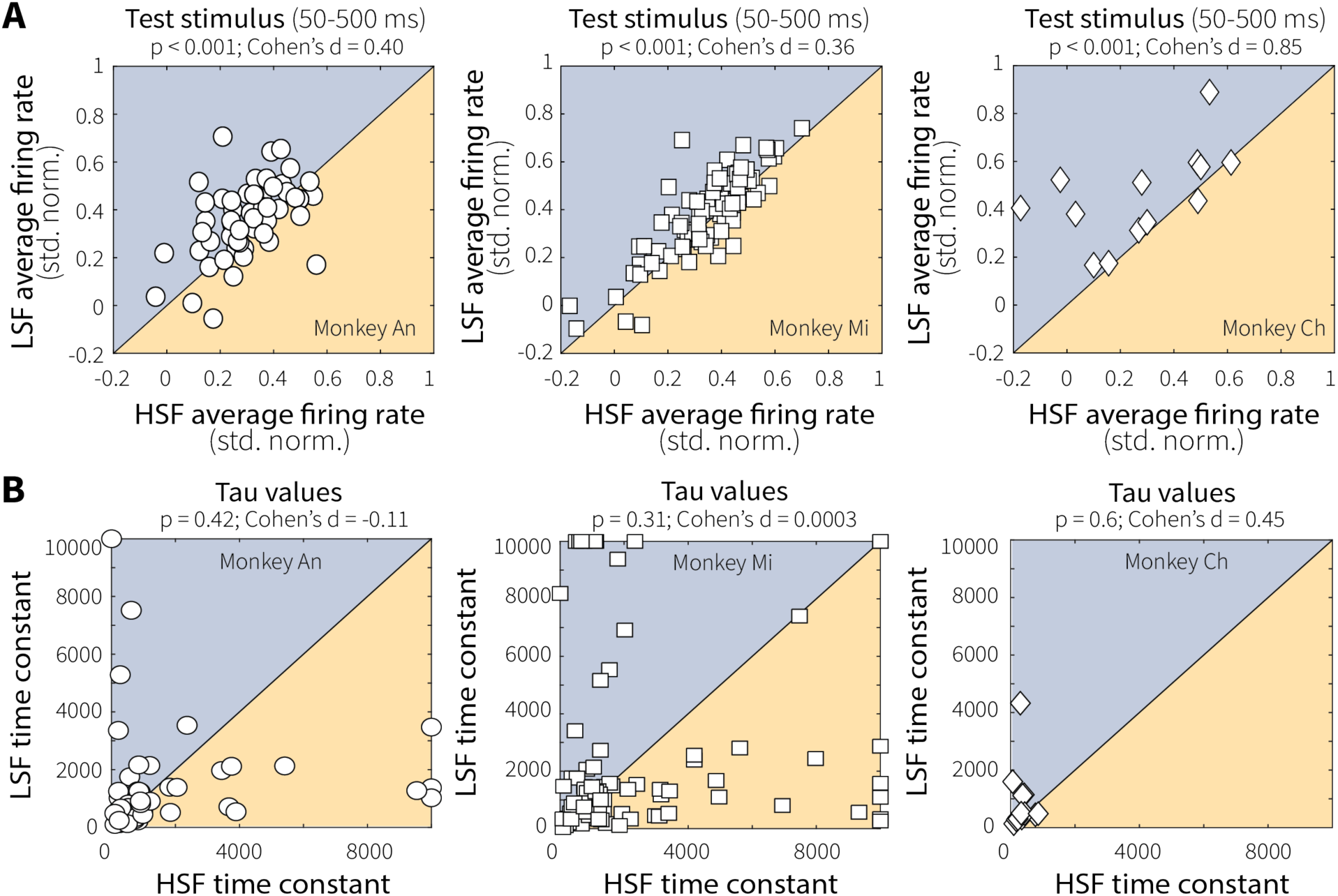
Context-dependent differences in evidence encoding were consistent across animals and reflected changes in response magnitude rather than temporal dynamics. (**A**) Pairwise comparisons of mean responses of individual MT units (points) to preferred motion during the test stimulus (50–500 ms after onset) for low (LSF) and high (HSF) switch-frequency conditions across sessions for each monkey, as indicated. All three monkeys exhibited stronger MT responses at LSF relative to HSF (*p*-values are from a Wilcoxon signed-rank test for equal medians), demonstrating consistent, context-stability-dependent differences in evidence encoding. (**B**) We quantified the time course of MT neural response by fitting baseline-subtracted, normalized activity during the test stimulus (preferred-motion switch trials only) with a single-exponential function. To facilitate fitting, data for each neuron were divided into three bins (200–330 ms, 340–470 ms, and 480–600 ms following test-stimulus onset) chosen to accommodate initial response latency and include approximately equal trial counts. Fitting was performed separately for low (LSF) and high (HSF) switch-frequency conditions. The exponential time constant (tau, in ms) from best-fitting, single-exponential fits showed no difference between LSF and HSF conditions for each monkey (*p*-values are from a Wilcoxon signed-rank test for equal medians). Thus, the temporal dynamics of MT responses were not reliably affected by recent temporal statistics.

**Extended Data Fig. 5:**
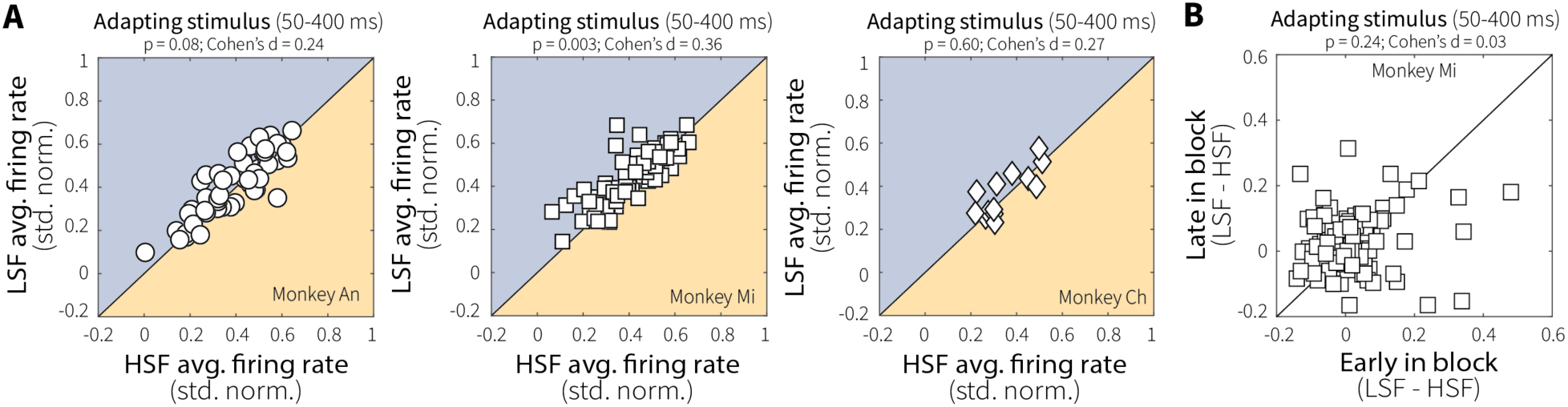
Context-dependent differences in initial MT responses were minimal and stable over the course of the session. (**A**) Average MT single-unit responses (points) during onset of the adapting stimulus (50–400 ms) for low (LSF) versus high (HSF) switch-frequency conditions, shown separately for each monkey. Initial responses to preferred motion were matched between LSF and HSF for Monkey An and Monkey Ch but were slightly elevated at LSF for Monkey Mi (*p*-values are from a Wilcoxon signed-rank test for equal medians). (**B**) For Monkey Mi, this initial difference in activity between LSF and HSF was stable across early and late blocks of each condition, possibly reflecting expectations about switching dynamics that were learned relatively quickly within each block of trials.

**Extended Data Fig. 6:**
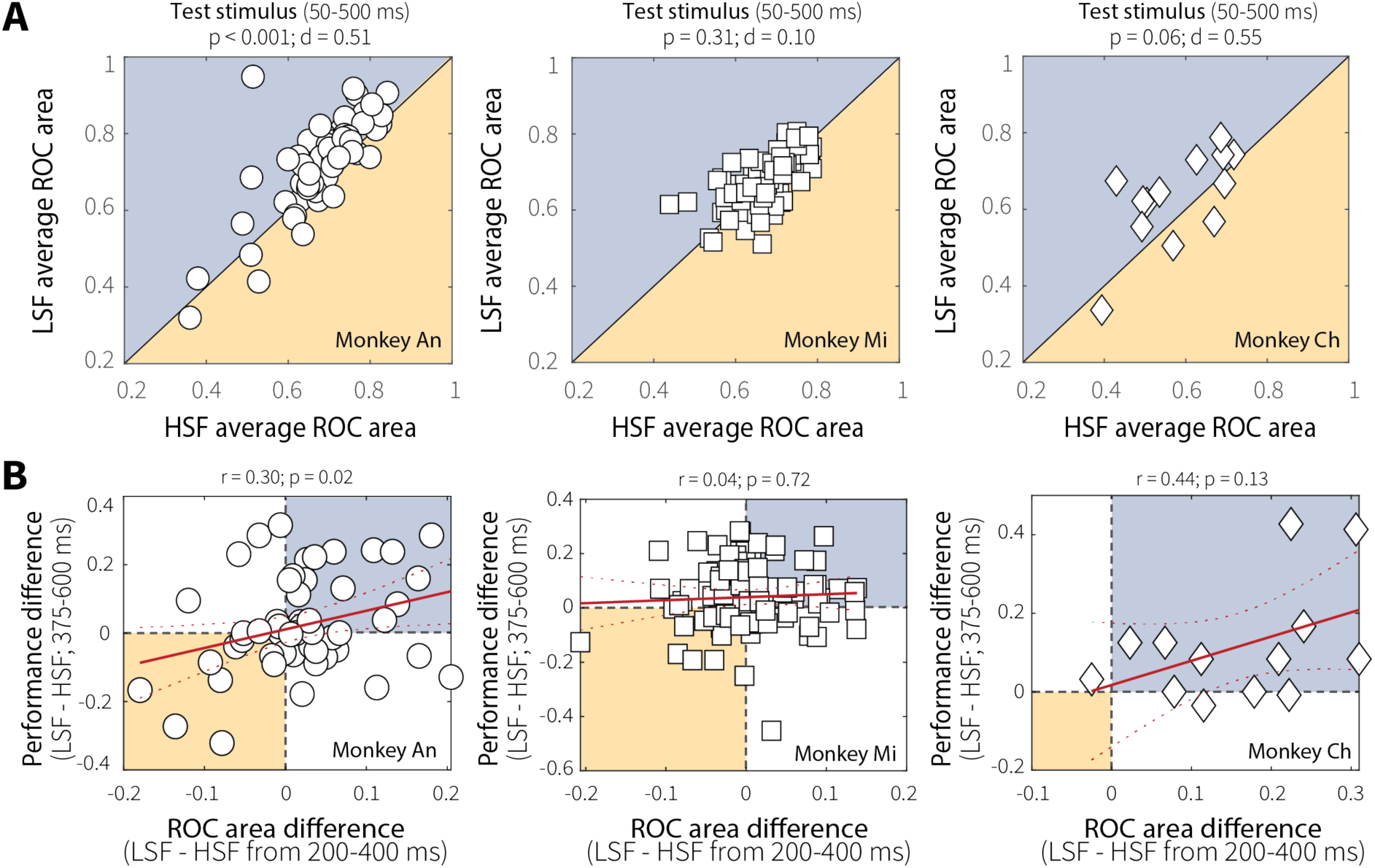
Context-dependent differences in MT evidence discriminability and its relationship to behavioral performance varied across monkeys. (**A**) Average ROC area during the stimulus (50–500 ms) for low (LSF) and high (HSF) switch-frequency conditions, shown separately for each monkey. ROC area was greater at LSF relative to HSF for Monkey An (Wilcoxon signed-rank test for equal medians). Monkey Ch showed a similar directional effect with a comparable effect size but for a much smaller sample and thus was not statistically significant. No effect was apparent for Monkey Mi despite a large sample size and context-stability differences in preferred-motion responses during the same stimulus window (Extended Data Fig. 4A). (**B**) Correlations between context-stability differences in ROC area (LSF–HSF; 200–400 ms) and behavioral performance (LSF–HSF percent correct; 375–600 ms), shown separately for each monkey. A significant positive relationship was observed for Monkey An, with a similar directional trend for Monkey Ch, though power was limited by the smaller number of recorded neurons. No relationship was observed for Monkey Mi despite a large sample size. Red solid and dashed lines indicate linear fits ± 95% CI.

**Extended Data Fig. 7:**
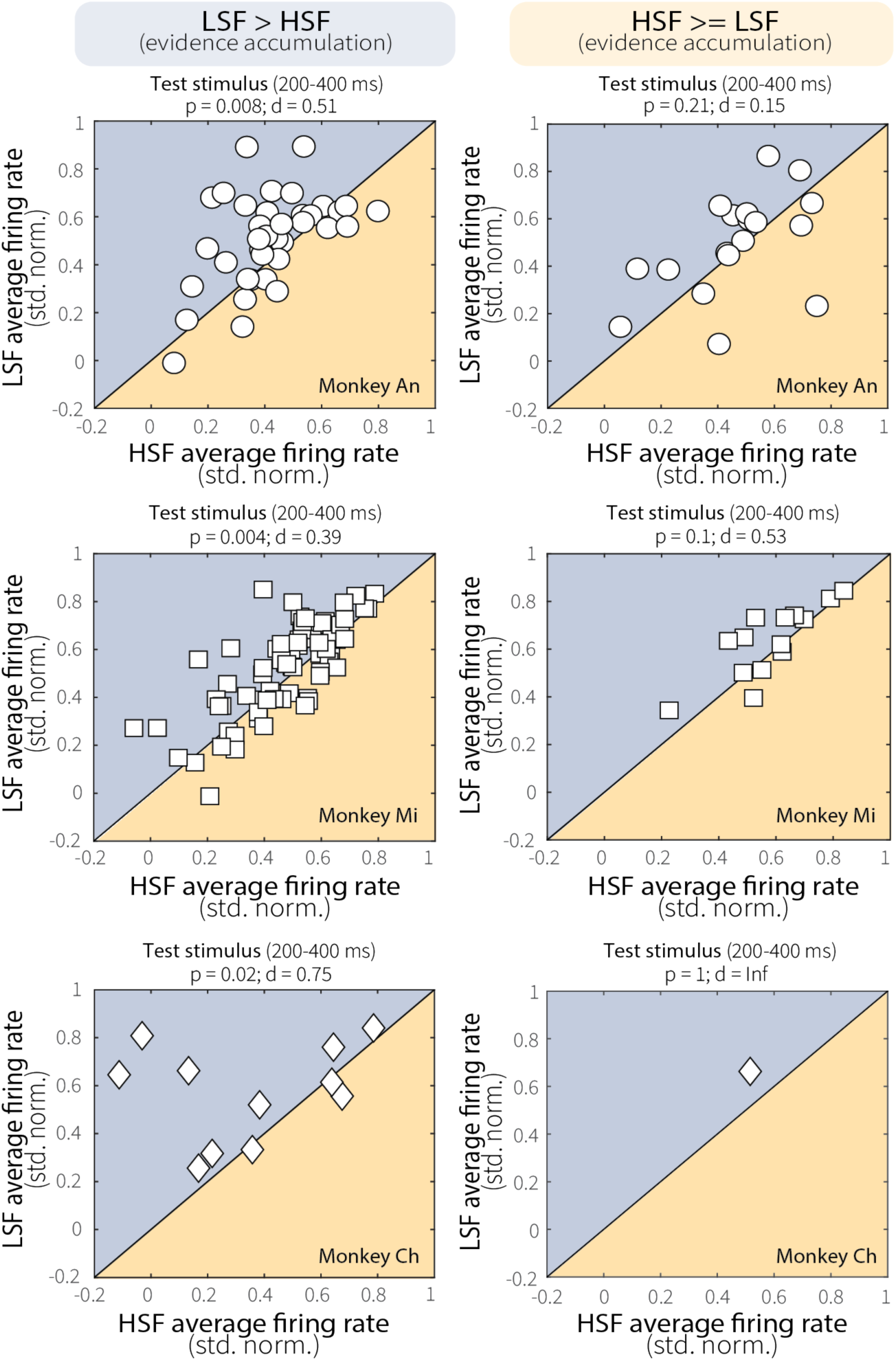
Relationship between MT neural activity and evidence-accumulation behavior was consistent across animals. Average MT neural activity during the test stimulus (200–400 ms) for low (LSF) and high (HSF) switch-frequency conditions, shown separately for each monkey (rows) and session group (columns). Sessions were divided based on evidence-accumulation behavior: *left column*: sessions in which psychometric slopes were steeper at LSF than HSF (LSF–HSF > 0); *right column*: sessions in which psychometric slopes were greater at HSF or equal across conditions (LSF–HSF ≤ 0). When the monkeys were more sensitive at LSF (left column), MT neurons showed greater activity at LSF relative to HSF. Conversely, when the monkeys were more sensitive at HSF or equally sensitive between conditions (right column), neural responses did not differ between conditions. *p*-values are for a Wilcoxon signed-rank tests for equal medians.

**Extended Data Fig. 8:**
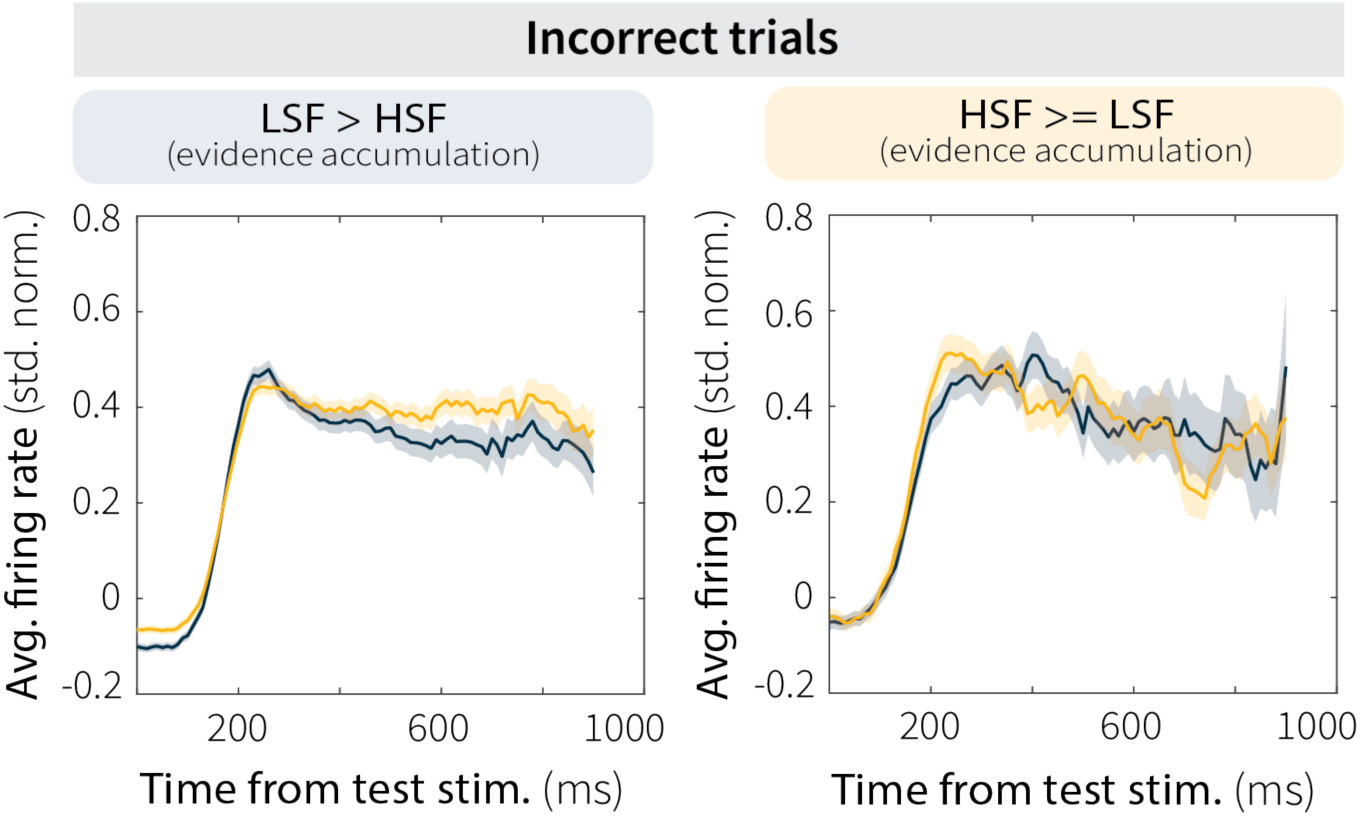
Context-dependent differences in MT activity were specific to correct trials. Baseline-subtracted and normalized MT firing rates during the test stimulus for low (LSF; blue) and high (HSF; orange) switch-frequency conditions, averaged across incorrect trials for all sessions. Data are shown separately for sessions in which the monkeys were more sensitive (i.e., accumulated more evidence over time) at LSF (left; LSF n = 1883, HSF n = 1939) or HSF (right; LSF n = 463, HSF n = 395). Unlike correct trials, error trials showed no consistent modulation by context stability or evidence-accumulation behavior, further confirming that context-dependent changes in MT neural activity were behaviorally relevant.

**Extended Data Fig. 9:**
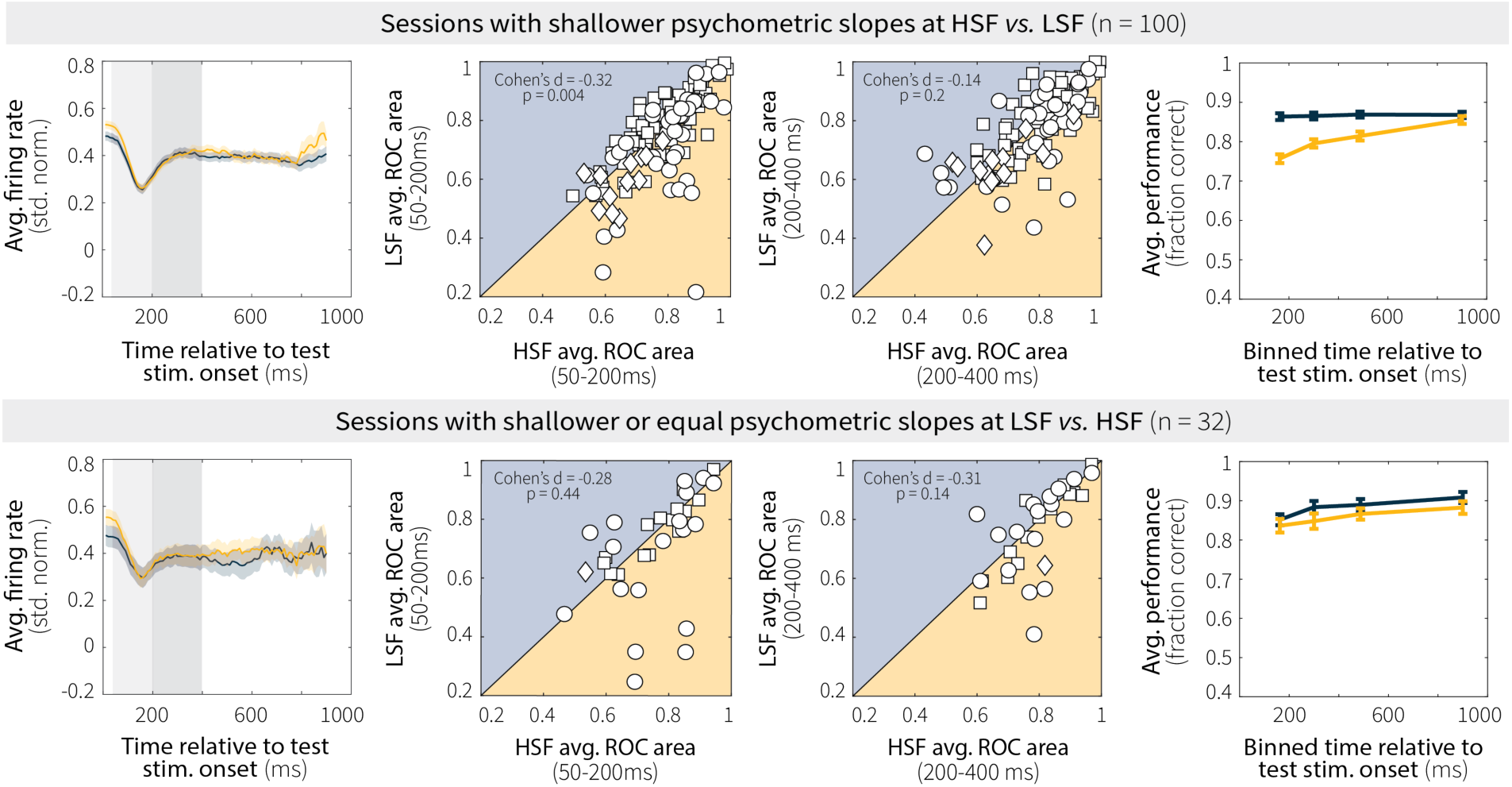
Context-dependent differences in MT neural activity and behavior on non-switch trials. Data are shown separately for sessions in which psychometric slopes were shallower at HSF *vs.* LSF (top row; n = 100) and psychometric slopes were shallower at LSF *vs.* HSF or equal across conditions (bottom row; n = 32). Columns show: (left) average MT firing rate during the test stimulus, (center-left and center-right) MT evidence discriminability (ROC area) in early (50–200 ms, lighter gray shading in the left panels) and late (200–400 ms, darker gray shading in the left panels) test-stimulus windows, and (right) behavioral performance as a function of test-stimulus duration. In the left panels, the transition from the adapting to the test stimulus, corresponding to a decrease in coherence, is visible as a decrease in MT neural activity ∼180 ms after test-stimulus onset. Thus, activity in the earlier time bin is likely driven partly by the adapting stimulus, whereas activity in the later time bin is likely driven mainly by the test stimulus. For sessions in which psychometric slopes were shallower at HSF vs. LSF (top row), there was a corresponding context-dependent difference in ROC area in the earlier (center-left), but not later (center-right), time bin. This difference in discriminability was not evident in the other sessions (bottom row, center-left). Consistent with this finding, context-dependent differences in behavioral performance on non-switch trials were also restricted to sessions with shallower psychometric slopes at HSF vs. LSF (top row, right panel). Moreover, in both session groups, behavioral performance on non-switch trials was near (or at) ceiling across viewing durations, reflecting the fact that monkeys accumulated evidence across adapting and test-stimulus epochs on non-switch trials. *p*-values are from Wilcoxon signed-rank tests for equal medians. Different symbols represent data from different monkeys (Monkey An, circles; Ch, diamonds; Mi, squares).

**Extended Data Fig. 10:**
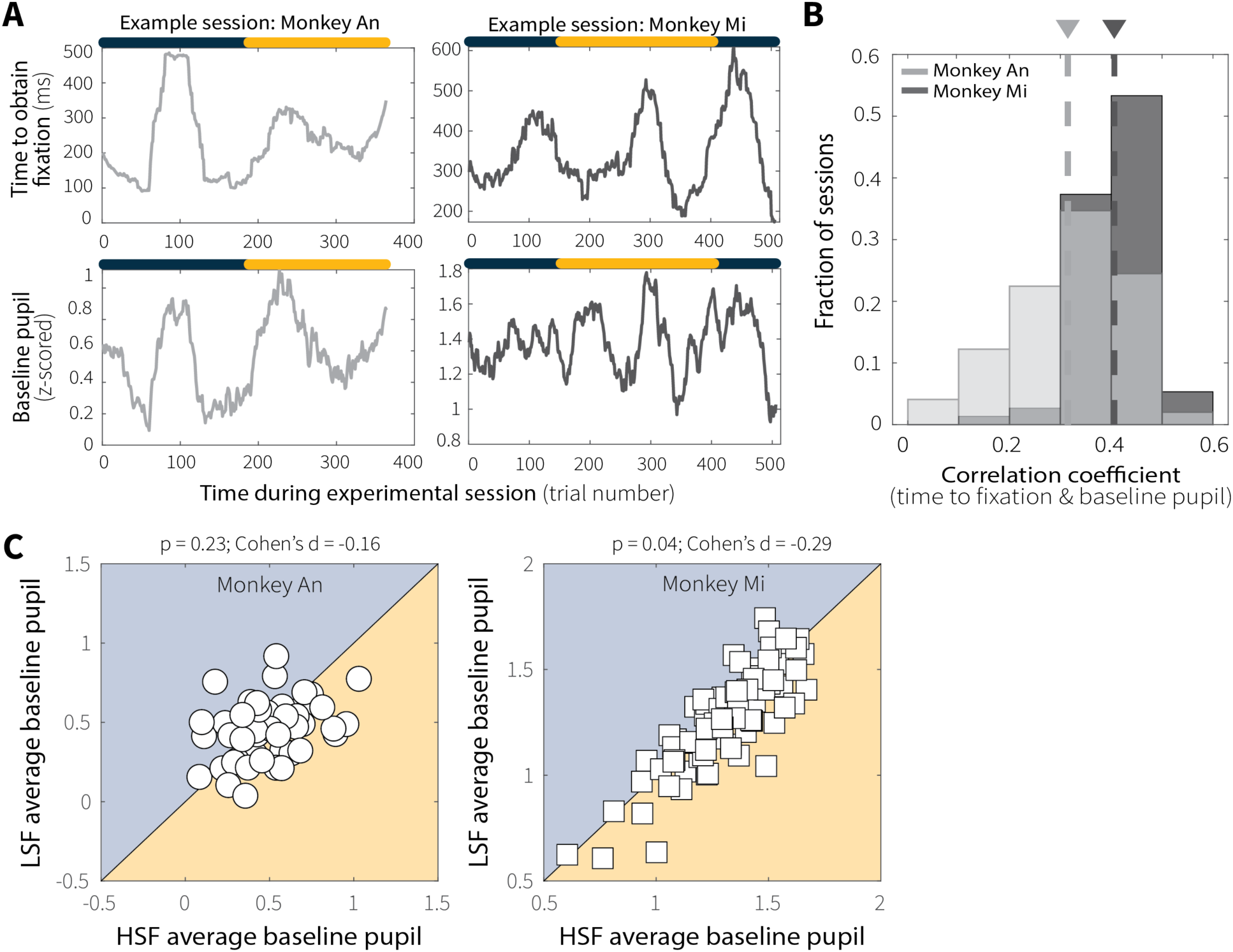
Baseline pupil diameter covaried with fixation-acquisition time across trials, sessions, and monkeys but did not differ across context-stability conditions. (**A**) Representative sessions illustrating trial-by-trial relationships between fixation-acquisition time (top) and baseline pupil diameter (bottom) for Monkey An (left, grey) and Monkey Mi (right, black). The colored bar above indicates the active context-stability condition on each trial. (**B**) Distributions of session-wise Spearman’s rank correlation coefficients between fixation-acquisition time and baseline pupil diameter across sessions for Monkey An (grey) and Monkey Mi (black). For both monkeys, the distribution of correlation coefficients was significantly positive (Monkey An: *p* < 0.001, t = 18.6, df = 48; Monkey Mi: *p* < 0.001, t = 48.93, df = 74). Mean correlation values are indicated by dashed lines and associated triangles. (**C**) Session-averaged baseline pupil diameter for LSF versus HSF conditions for Monkey An (left) and Monkey Mi (right). Average baseline pupil size did not reliably differ across context-stability conditions for monkey An and only slightly for monkey Mi (*p* values are for a Wilcoxon signed-rank test for equal medians).

**Extended Data Fig. 11:**
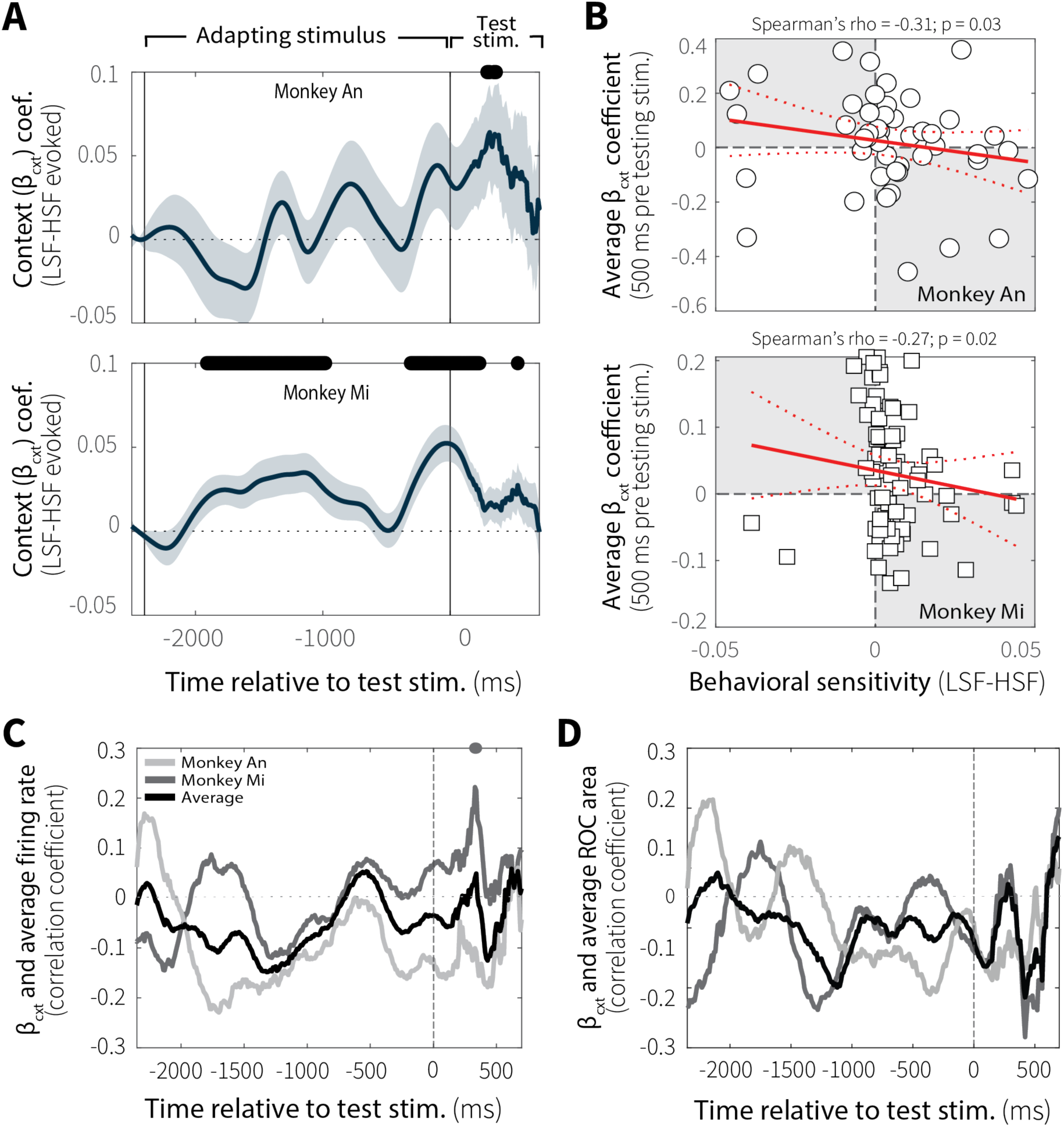
Evoked pupil diameter related to context-dependent evidence accumulation-behavior, but not MT neural activity. (**A**) Context-stability effects on evoked pupil diameter for individual monkeys. Sliding-window regression coefficients (β_cxt_; mean ± SEM across sessions) estimating differences in pupil diameter between low (LSF) and high (HSF) switch-frequency conditions while controlling for baseline pupil size. Black bar (top) indicates windows in which β_cxt_ differed significantly from zero (*p* < 0.05, uncorrected for multiple comparisons). (**B**) Average evoked pupil differences (β₂, −500–0 ms relative to test-stimulus onset) plotted versus behavioral sensitivity differences (LSF–HSF psychometric slope) from individual sessions for Monkey An (top) and Monkey Mi (bottom). Red solid and dashed lines indicate a linear fit ± 95% CI. (**C**) Spearman’s rank correlation between context-stability differences in MT neural activity (LSF–HSF, 50–500 ms) and pupil β_cxt_ coefficients from individual sessions as a function of time relative to test-stimulus onset (computed in 100 ms bins with a 10 ms slide). Individual animals are shown separately (Monkey An: light grey; Monkey Mi: dark grey) along with the across-animal average (black). Corresponding colored bars indicate time points with significant correlations (*p* < 0.05, uncorrected for multiple comparisons). (**D**) Same analysis for MT discriminability (LSF–HSF ROC area, 50–500 ms) and pupil β_cxt_ coefficients as a function of time relative to test-stimulus.

**Extended Data Fig. 12:**
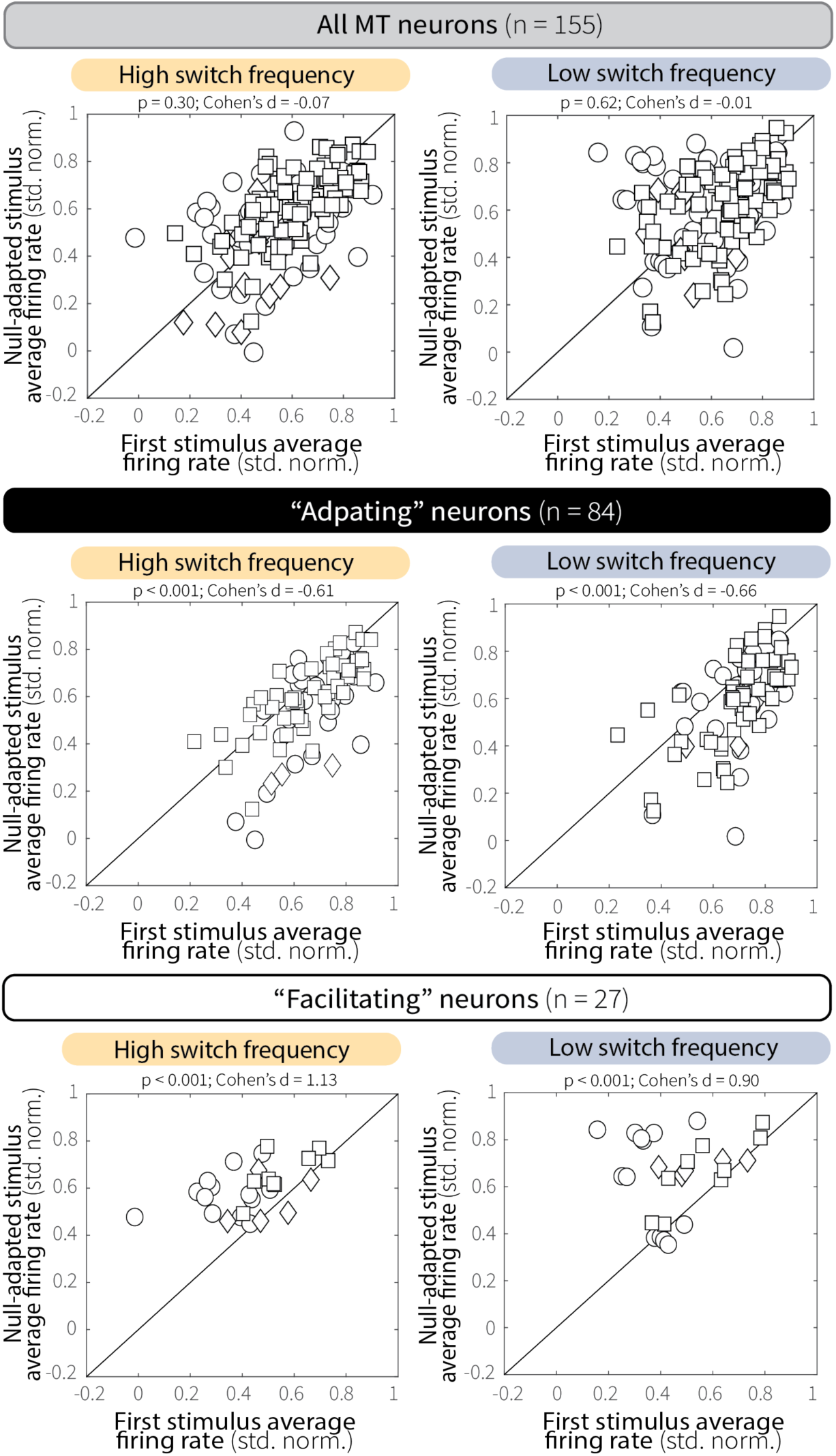
Effects of null-motion adaptation on preferred-motion responses in MT. Pairwise comparisons of MT responses to preferred motion before (first stimulus; solid orange line in Fig. 3C) versus after null motion (second, null-adapted stimulus; dashed orange line in Fig. 3C), shown separately for high (HSF, left) and low (LSF, right) switch-frequency conditions. Across the population (top row), prior exposure to null motion did not reliably alter subsequent preferred-motion responses. However, null-motion adaptation produced opposite effects in subgroups of adapting and facilitating neurons. Adapting single units (middle row) showed modest decreases in response to preferred motion following null-motion exposure, whereas facilitating units (bottom row) showed increased responses. Notably, the reduction observed in adapting single units was smaller than that following repeated preferred-motion stimulation. p-values are from Wilcoxon signed-rank tests for equal medians.

